# Loss of PINK1 Causes Age-dependent Mitochondrial Trafficking Deficits in Nigrostriatal Dopaminergic Neurons Through Aberrant Activation of p38 MAPK

**DOI:** 10.1101/2024.07.17.602426

**Authors:** Jingyu Zhao, Yuanxin Chen, Lianteng Zhi, Qing Xu, Hui Zhang, Chenjian Li

## Abstract

Mutations in *PTEN-induced putative kinase 1* (*PINK1*) cause early-onset, autosomal-recessive Parkinson’s disease (PD). While previous studies showed age-related declines in dopamine release and ATP levels in *Pink1^-/-^* mice, the mechanisms remain unclear. Using a novel TH-Mito-Dendra2 transgenic mouse model to label dopaminergic neuron mitochondria, it is found that PINK1 loss leads to age-dependent deficits in mitochondrial axonal trafficking, characterized by reduced anterograde movement and increased static mitochondria in acute brain slices, which more closely mimic *in vivo* conditions. Pharmacological induction of reactive oxygen species (ROS) and calcium release impaired mitochondrial mobility. *Pink1* knockout mice exhibited elevated mitochondrial calcium, ROS levels, and p38 MAPK hyperactivation. Treatment with p38 inhibitor SB202190 restored mitochondrial motility and increased anterograde transport. Our findings suggest that PINK1 loss disrupts mitochondrial trafficking by disturbing calcium homeostasis and ROS production via the p38 pathway, contributing to PD pathogenesis.

**Table of Contents:** 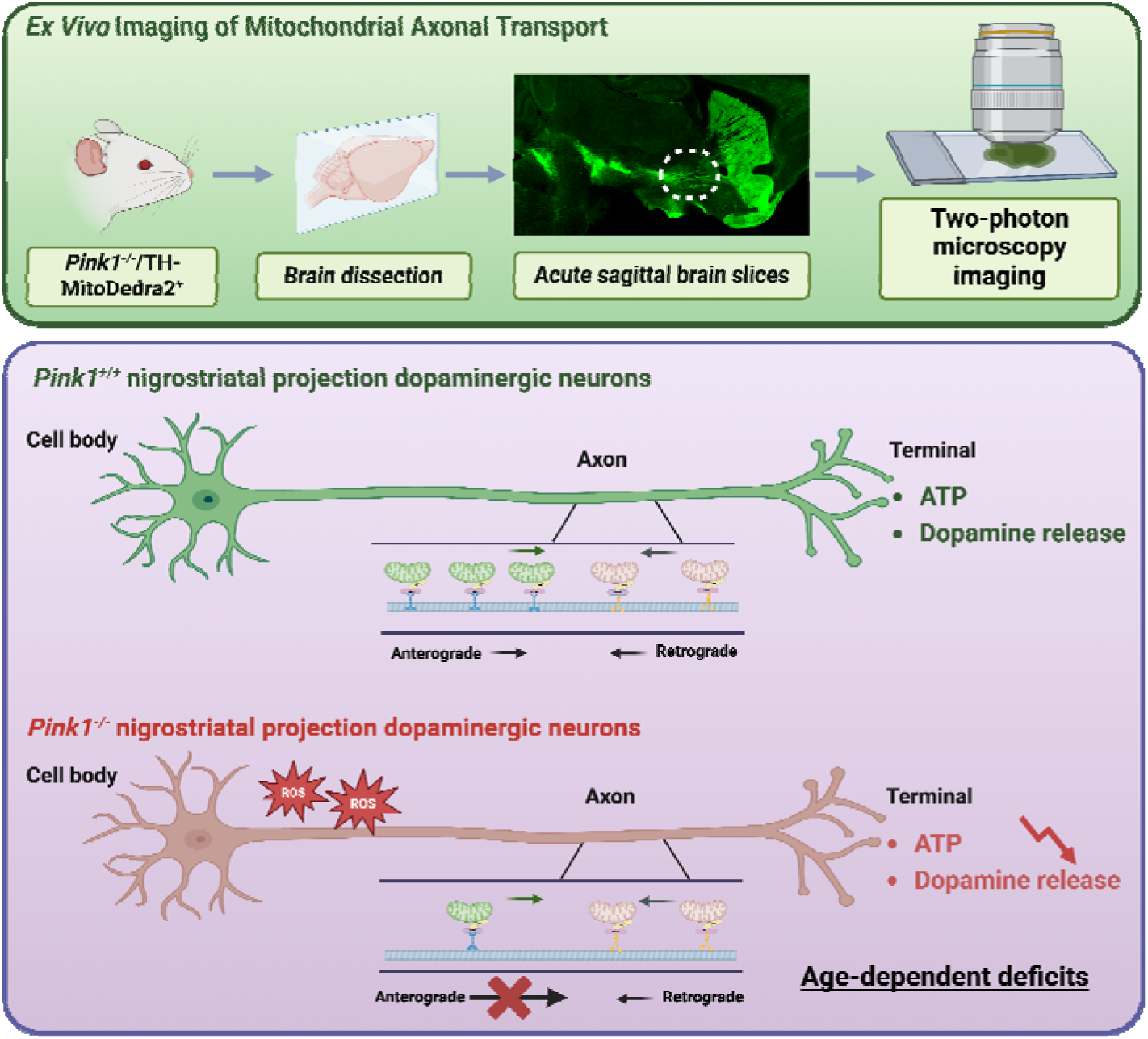

Using a novel mouse model with labeled dopaminergic neuron mitochondria, Zhao J. et al. discovered age dependent disruptions in mitochondrial transport. Their research showed progressive deterioration of mitochondrial movement, characterized by reduced anterograde movement and increased static mitochondria. These changes were associated with higher oxidative stress and hyperactivation of p38 MAPK in Parkinson’s disease.

## Introduction

Parkinson’s disease (PD) is a major neurodegenerative disorder affecting people above 65 years old, characterized by dopamine neuron loss in the substantia nigra pars compacta (SNc) leading to striatal dopamine deficiency^1^. While the pathogenesis of PD is complicated, mounting evidence implicates mitochondrial dysfunction as a key initiating factor^2^. Mitochondrial abnormalities in PD include altered morphology, impaired energy production, increased oxidative stress, dysregulated dynamics and calcium homeostasis, and disrupted mitophagy. However, it remains unclear whether these mitochondrial defects are primary causes or downstream consequences in PD. The identification of *PINK1* as a PD gene provided direct evidence linking mitochondrial dysfunction to PD pathogenesis^3^. PINK1 regulates key mitochondrial processes like mitophagy, calcium homeostasis, and fission in neurons, underscoring its relevance in studying mitochondrial dysfunction’s role in PD. PINK1 loss-of-function models demonstrate mitochondrial defects like increased fission^4^, depolarization^5^, impaired electron transport chain complexes I and III^6,7^, and calcium overload^8^. Additionally, PINK1 interacts with Miro/Milton, implicating its role in mitochondrial trafficking^9^.

Neurons, especially synapses, are highly energy demanding. A human cortical neuron utilizes ∼4.7 billion ATPs/second at rest^10^, underscoring the importance of mitochondrial trafficking. Following evidence that PINK1 can interact with trafficking adaptor proteins^9^, several studies have explored the relationship between PINK1 and mitochondrial trafficking in different models. One study demonstrated that overexpression of PINK1 or PARKIN in rat hippocampal axons significantly reduced mitochondrial motility. For impaired mitochondria, PINK1 phosphorylates Miro, leading to PARKIN-dependent Miro degradation and subsequent dissociation of kinesin from mitochondria, thus sequestering depolarized mitochondria^11^. Another study demonstrated that D-AKAP1/PKA (D-AKAP1, dual-specificity A Kinase Anchoring protein 1; PKA, Protein Kinase A) can restore the mitochondrial trafficking deficiency in cortical neurons from PINK1 knockout (KO) mice^12^. While these studies support the critical role of PINK1/PARKIN pathway in regulating mitochondrial trafficking^13^, most research utilizes *in vitro* systems or *Drosophila* models, which differ from *in vivo* conditions. Additionally, most studies are not specific to dopaminergic neurons. From this perspective, further investigations closer to *in vivo* settings and focused on dopaminergic neurons are needed.

Here, we generated a new Mito-Dendra2 tool mouse model to specifically label mitochondria in dopaminergic neurons and established a new approach to directly observe mitochondrial trafficking in nigrostriatal dopaminergic neurons in acute brain slices from a *Pink1^-/-^* mouse model. We found an age-dependent impairment of mitochondrial motility and anterograde transport in dopaminergic neurons lacking PINK1 mediated through the aberrant p38 pathway. These results provide a possible explanation for our previous findings of decreased dopamine release and ATP production in the striatum of *Pink1^-/-^* mice at early middle-age^14^.

## Results

### Generation and characterization of TH-MitoDendra2 BAC transgenic mice

To specifically label mitochondria in dopaminergic neurons, we generated a bacterial artificial chromosome (BAC) clone encompassing the entire tyrosine hydroxylase (*Th*) gene (Fig. 1 A). A mitochondria-targeting photo-convertible fluorescent protein Dendra2 gene^15,16^ was inserted in the *Th* exon 1 (Fig. 1 A). The co-localization of Mito-Dendra2 and the immunofluorescence for TH confirmed the specificity of Mito-Dendra2 in dopaminergic neurons within the SNc and ventral tegmental (VTA) area (Fig. 1 B-E). We also verified the specificity of Mito-Dendra2 signals at the dopaminergic terminals in the striatum and olfactory bulb (Fig. S1) and the results showed Mito-Dendra2 only expressed in TH positive neurons and neurites. Moreover, the co-localization of Mito-Dendra2 signals with MitoTracker staining verified the subcellular localization of Mito-Dentra2 in mitochondria (Fig. 1 F). Using this model, we can visualize the continuous and intact projection bundles from dopaminergic neurons in the SNc to the striatum in the sagittal view (Fig. 1 E, S2). These results demonstrate that Mito-Dendra2 is specifically localized in the mitochondria of dopaminergic neurons and their projections, providing a novel and powerful tool for studying mitochondrial trafficking in these neurons.

**Fig. 1.**
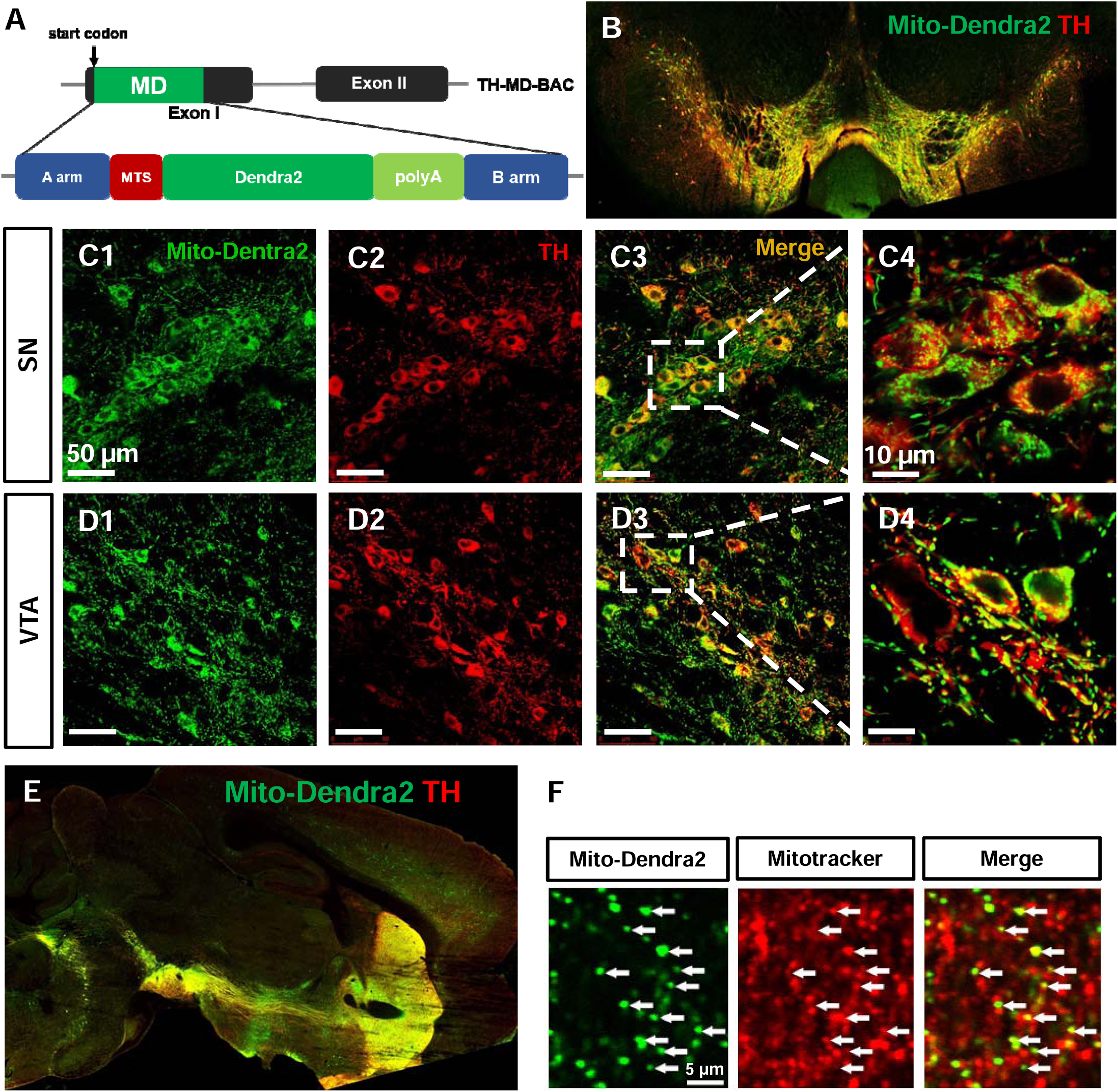
Generation and characterization of TH-Mito-Dendra2 (MD) BAC transgenic mouse. **(A)** Depiction of the TH-MitoDendra2 BAC construct. To modify RPCI23-350E13 BAC, a targeting cassette was generated with A arm, B arm and mitochondrial-targeting dendra2 insert. The mito-dentra2 and Th gene exon1 share the same initiation ATG codon. A and B represent the homologous recombination regions. MTS: mitochondrial targeting sequence. **(B)** A coronal section of TH-Mito-Dendra2 (MD) BAC transgenic mouse brain showing the histological overview of Mito-Dentra2 expression pattern. Tyrosine hydroxylase (TH) signal (red) co-localizes with Mito-Dentra2 fluorescence (green) in midbrain areas. **(C)** Mito-Dendra2 (green) expressing cells are all TH (red) immunoreactive in the substantia nigra (SN). **(D)** Mito-Dendra2 (green) expressing cells are all TH (red) immunoreactive in the ventral tegmental area (VTA). **(E)** A sagittal section of TH-MitoDendra2 BAC mouse brain showing continuous and intact nigrostriatal axonal projection of dopaminergic neurons. **(F)** MitoDendra2 is localized in mitochondria, as labeled with MitoTracker, in the striatum of TH-mitodendra2-BAC transgenic mice.

### Investigation of mitochondrial trafficking in acute brain slices with two-photon laser microscopy (2PLSM)

To evaluate mitochondrial trafficking in *Pink1^-/-^* mice, we crossed them with Mito-Dendra2 transgenic mice to generate *Pink1^-/-^* /MD+ transgenic mice (Fig. 2 A). We chose to use acute sagittal brain slices (Fig. 2 A), which closely mimic *in vivo* conditions, to monitor mitochondrial trafficking. Two-photon laser scanning microscopy (2PLSM) was employed to track mitochondrial movement. Using time-lapse imaging, we visualized and recorded videos showing mitochondrial trafficking along the axon bundles (Fig. 1 E, Fig. 2 A, and B). Mitochondrial movement was primarily classified into three states: stationary, anterograde (moving towards the terminals), and retrograde (moving towards the cell body). As depicted in Figure 2 B and Video S1, we clearly captured mitochondrial movement at sequential time points. After collecting mitochondrial trafficking data, we used the software Fiji with two plugins, KymographDirect and KymographClear^17^, to analyze the differences in the percentage and velocity of moving mitochondria (Fig. 2 C).

**Fig. 2.**
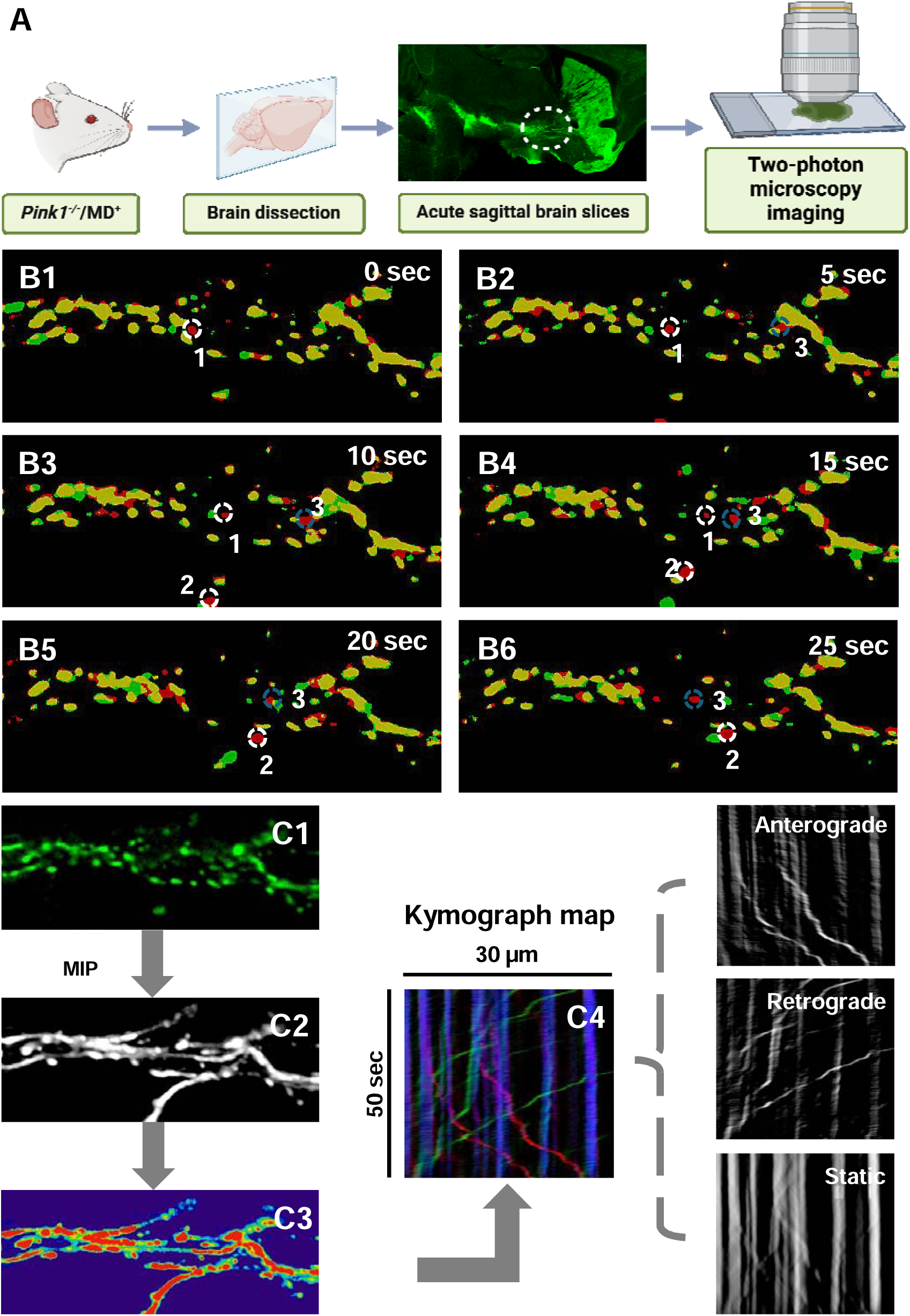
Investigation of mitochondrial movement in acute brain slices using two-photon laser microscopy (2PLSM). **(A)** Schematic illustration of mitochondrial trafficking imaging in acute brain slices. **(B)** Image sequences of mitochondrial trafficking. B1-B6 represent images of moving mitochondria at the different time points: 0, 5, 10, 15, 20 and 25 seconds, respectively. White circles 1 and 2 denote the anterograde movement of mitochondria, blue circle 3 denotes the retrograde movement of mitochondria. **(C)** Kymograph generation and mitochondria trafficking measurement. C1: Representative image of mitochondria expressing Dendra2 protein. C2: Maximum intensity projection of a series of mitochondrial trafficking images. C3: Pseudo color map of panel C2 to facilitate outlining the trace of mitochondrial movement. C4: Kymograph map generated from the images, showing anterograde, retrograde, and static traces. Red lines indicate the anterograde trace of mitochondrial movement, green lines indicate the retrograde movement, and blue lines represent static mitochondria.

### Age-dependent mitochondrial axonal transport deficits in *Pink1^-/-^* mice

After establishing the procedure, we evaluated the alterations in mitochondrial trafficking between *Pink1^-/-^* mice and age-matched wild-type (*Pink1^+/+^*) littermates at different ages (Fig. 3, Tables S1-S3). The percentages of stationary, anterograde, and retrograde mitochondria were similar in 2-4-month-old *Pink1^-/-^* mice and WT littermates (Fig. 3 A1-A2, B1-B2, C, Table S1). The average velocities of anterograde and retrograde movement were also comparable (Fig. 3 D, Table S1). Approximately 84-85% of mitochondria were stationary in both genotypes at 2-4 months, which is higher than typically reported in cultured cells^13,18^. In these mice, about 11% of mitochondria moved anterogradely and 4-5% moved retrogradely, differing from the more balanced 15% anterograde and 15% retrograde movement seen in cultured cells^19^.

**Fig. 3.**
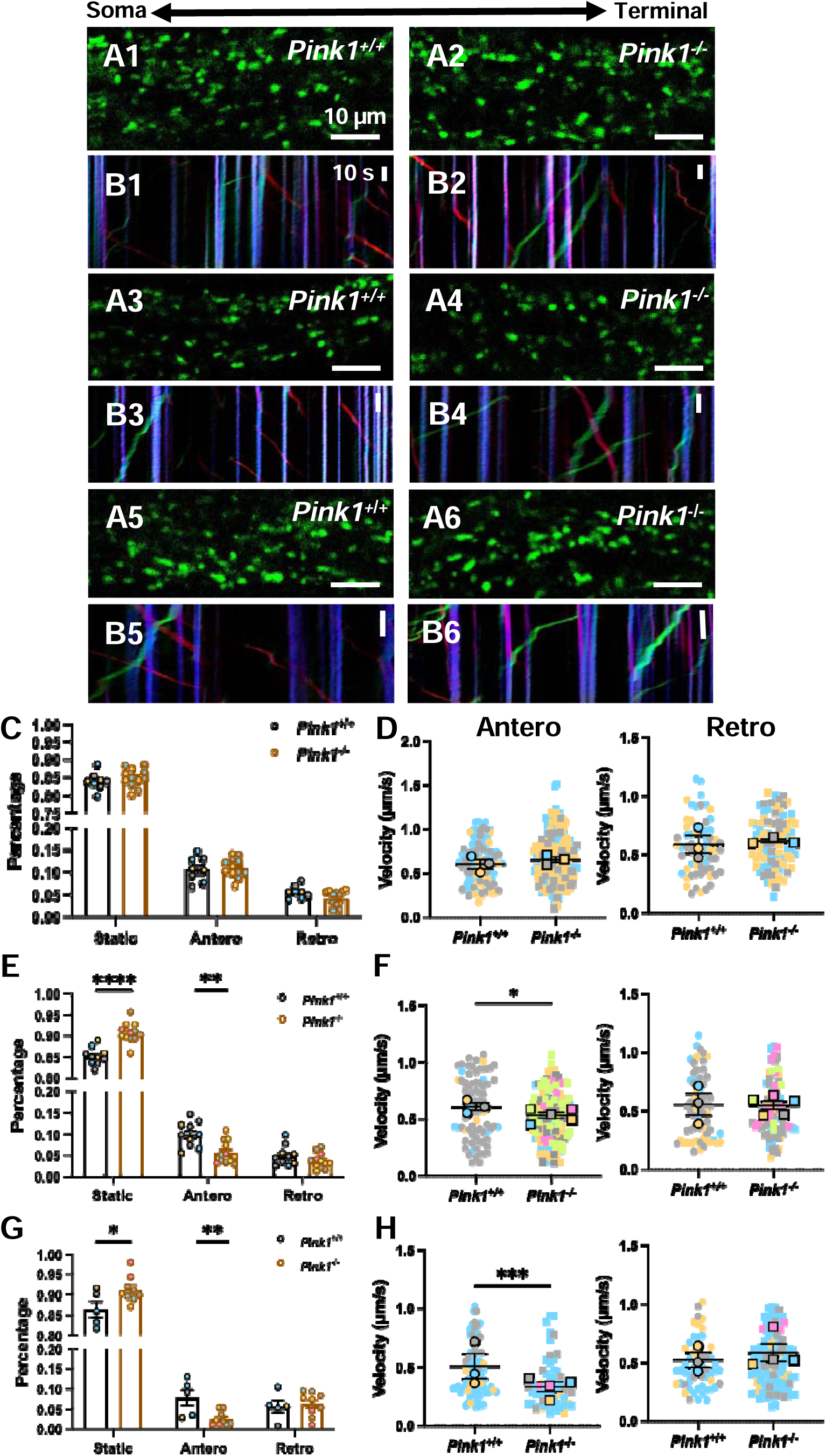
Age-dependent mitochondrial axonal transport deficits in *Pink1^-/-^* mice. **(A)** Representative *in vivo* two-photon microscopy images of mitochondrial trafficking in dopaminergic neuron axons at the initial time point. Images depict *Pink1^+/+^* and *Pink1^-/-^* mice at 2-4 months (A1-A2), 5-7 months (A3-A4), and above 10 months (A5-A6) of age, respectively. **(B)** Representative kymographs illustrating mitochondrial trafficking patterns in *Pink1^+/+^* and *Pink1^-/-^* mice across different age groups. **(C)** Quantification of mitochondrial motility states in 2-4-month-old *Pink1^+/+^* (N = 3, n = 8) and *Pink1^-/-^* (N = 3, n = 12) mice. Data points represent mean percentages per brain slice, with colors denoting individual animals. *P*-values were determined using the unpaired t-test. **(D)** SuperPlots illustrating anterograde (left) and retrograde (right) mitochondrial velocities in 2-4-month-old mice. In scatter plots, each data point represents an individual mitochondrial velocity measurement. In summary plot, each data point represents the mean velocity per animal, with colors distinguishing individual mice. *P*-values were determined using the Mann-Whitney U test. **(E)** Quantification of mitochondrial motility states in 5-7-month-old *Pink1*^+/+^ (N = 3, n = 10) and *Pink1*^-/-^ (N = 5, n = 12) mice. *P*-values were determined using the unpaired t-test. \*\**P* < 0.01, \*\*\*\**P* < 0.0001. **(F)** SuperPlots illustrating anterograde (left) and retrograde (right) mitochondrial velocities in 5-7-month-old mice. *P-*values were determined using the Mann-Whitney U test. \**P* < 0.05. **(G)** Quantification of mitochondrial motility states in older than 10-month-old *Pink1*^+/+^ (N = 3, n = 5) and *Pink1*^-/-^ (N = 4, n = 9) mice. *P*-values were determined using the unpaired t-test. \**P* < 0.05, ** *P* < 0.01. **(H)** SuperPlots illustrating anterograde (left) and retrograde (right) mitochondrial velocities in mice older than 10 months. *P*-values were determined using the Mann-Whitney U test. \*\*\**P* < 0.001.

In 5-7-month-old *Pink1*^-/-^ mice, the proportion of stationary mitochondria increased significantly (90.79 ± 0.69% (*Pink1*^-/-^) *vs.* 85.18 ± 0.65% (*Pink1^+/+^*), *P* < 0.0001). Anterograde movement decreased from 9.95 ± 0.94% to 5.63 ± 0.77% (*P* = 0.0019), while retrograde movement showed a slight decrease (3.58 ± 0.52% (*Pink1*^-/-^) *vs.* 4.88 ± 0.75% (*Pink1^+/+^*), *P* = 0.1624; Figure 3 A3-A4, B3-B4, E, Table S2). Additionally, the anterograde velocity decreased significantly (0.52 ± 0.02 µm/s (*Pink1*^-/-^) *vs.* 0.60 ± 0.03 µm/s (*Pink1^+/+^*), *P* = 0.0335) (Fig. 3 F, Table S2). This reduction in anterograde movement may result in less mitochondrial content in axonal terminals.

In *Pink1*^-/-^ mice older than 10 months, the percentage of static mitochondria increased significantly compared to WT littermates (91.72 ± 1.07% (*Pink1*^-/-^) *vs.* 86.55 ± 1.88% (*Pink1^+/+^*), *P* = 0.0251), while anterograde movement decreased to 2.54 ± 0.60% (*P* = 0.008) (Fig. 3 A5-A6, B5-B6, G, Table S3). In these older *Pink1*^-/-^ mice, retrograde mitochondrial transport exceeded anterograde transport, reaching 6.24 ± 1.00% (Fig. 3 G, Table S3). Furthermore, the anterograde velocity was significantly decreased to 0.36 ± 0.03 µm/s (*P* = 0.004) compared to age-matched WT controls (0.48 ± 0.02 µm/s), while retrograde velocity remained unchanged (Fig. 3 H, Table S3).

In summary, *Pink1*^-/-^ mice exhibited age-dependent deficits in mitochondrial axonal trafficking. These deficits, which were not present in 2-4-month-old mice, emerged around 5 months of age and persisted beyond 10 months of age. The trafficking abnormalities included an increased proportion of static mitochondria, a decreased proportion of antegrade motile mitochondria, and slower anterograde velocities. These findings differ from previous reports in cultured neurons^11^ and *Drosophila*^20^, highlighting potential differences between *in vivo* and *in vitro* models, as well as between invertebrate and vertebrate systems.

### Mitochondrial calcium elevation and transport deficits in *Pink1^-/-^* mice

Mitochondrial axonal transport is a tightly regulated process involving many factors. Cytosolic Ca^2+^ can modulate mitochondrial motility via the Miro protein^21^. Furthermore, mitochondria themselves play a pivotal role in calcium homeostasis^22^. We first evaluated mitochondrial calcium levels in 4-month-old *Pink1*^-/-^ mice and WT littermates using AAV9-TH-mito-GCaMP6, a modified GCaMP6 virus that is specifically expressed in mitochondria in dopaminergic neurons. Figure 4 A and B show that the relative [Ca^2+^]_mt_ (relative fluorescence intensity) was significantly increased in *Pink1*^-/-^ mice (78.6 ± 4.3%) compared to WT littermates (35.7 ± 4.6%, *P* = 0.0001). Next, we investigated the effect of elevated mitochondrial calcium on mitochondrial trafficking. We incubated WT acute brain slices with 5 µM CGP37157, an inhibitor of the mitochondrial Na⁺/Ca²⁺ exchanger (NCLX) that causes mitochondrial calcium overload^23^. Mitochondrial motility was significantly decreased after 2 hours of CGP37157 incubation (Fig. 4 C-D, Video S2 and S3). The proportion of static mitochondria increased from 85.26 ± 0.69% to 90.29 ± 1.17% (*P* = 0.0048), while anterograde movement declined from 8.69 ± 1.07% to 5.62 ± 0.85% (*P* = 0.0432) (Fig. 4 D).

**Fig. 4.**
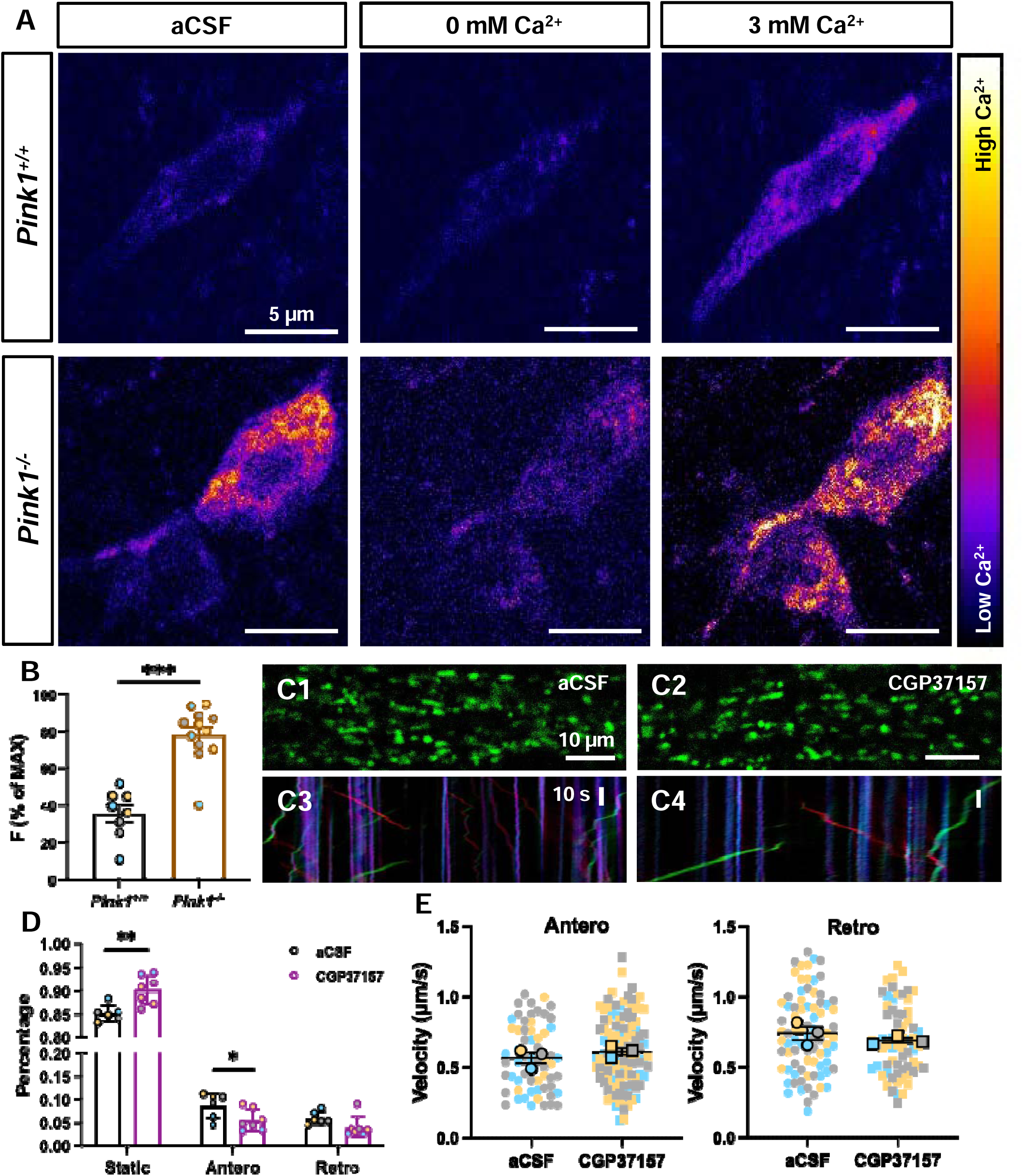
Mitochondrial calcium elevation and transport deficits in acute brain slices. **(A)** Representative fluorescence images of TH-mito-GCaMP6 in acute brain slices from approximately 3-month-old *Pink1*^+/+^ (upper panel) and *Pink1*^-/-^ mice (lower panel). Images depict mitochondrial [Ca^2+^] under baseline conditions (left column), after 20 minutes of Ca^2+^-free aCSF perfusion (500 µM EGTA; middle column) and following 30 minutes of 3 mM Ca^2+^ aCSF perfusion (right column). **(B)** Quantification of mitochondrial [Ca^2+^] in *Pink1^+^*^/+^ (N = 3, n = 8) and *Pink1*^-/-^ (N = 3, n = 12) mice. Data points represent mean fluorescence intensity per brain slice, with colors denoting individual animals. Statistical significance was determined using the Mann-Whitney U test. \*\*\**P* = 0.0001. **(C)** Representative images and corresponding kymographs illustrating mitochondrial dynamics before (C1 and C3) and after (C2 and C4) 2-hour incubation with 5 µM CGP37157. **(D)** Quantification of mitochondrial motility states following 5 µM CGP37157 treatment. Ctrl: N = 3, n = 6; CGP37157: N = 3, n = 7. *P*-values were determined using the unpaired t-test. \**P* < 0.05, \*\**P* < 0.01. **(E)** SuperPlots of anterograde (left) and retrograde (right) mitochondrial velocities in control and CGP37157-treated groups. *P*-values were determined using the Mann-Whitney U test.

This phenomenon of altered mitochondrial motility following [Ca^2+^]_mt_ elevation was reminiscent of the findings observed in 5- to 7-month-old and above 10-month-old *Pink1*^-/-^ mice, suggesting a similar underlying mechanism contributing to the mitochondrial trafficking defects in *Pink1*^-/-^ mice. The velocities of anterograde and retrograde mitochondrial transport remained unaffected by the calcium perturbations in this experimental context (Fig. 4 E).

### Reduced mitochondrial anterograde movement by ROS elevation

Mitochondrial calcium overload impairs normal mitochondrial function, leading to increased reactive oxygen species (ROS) generation and decreased ATP production^24,25^. To elucidate the relationship between mitochondrial calcium homeostasis and reactive oxygen species (ROS) generation in acute brain slices, we utilized CGP37157 to induce mitochondrial calcium overload in brain slices derived from roEGFP transgenic mice (4-5-month-old) and evaluate the ROS alteration^26^ after treatment (Fig. 5). Our results showed a significant increase in [Ca^2+^]_mt_, which led to an elevation in ROS levels (*P* = 0.0005, Fig. 5 C).

**Fig. 5.**
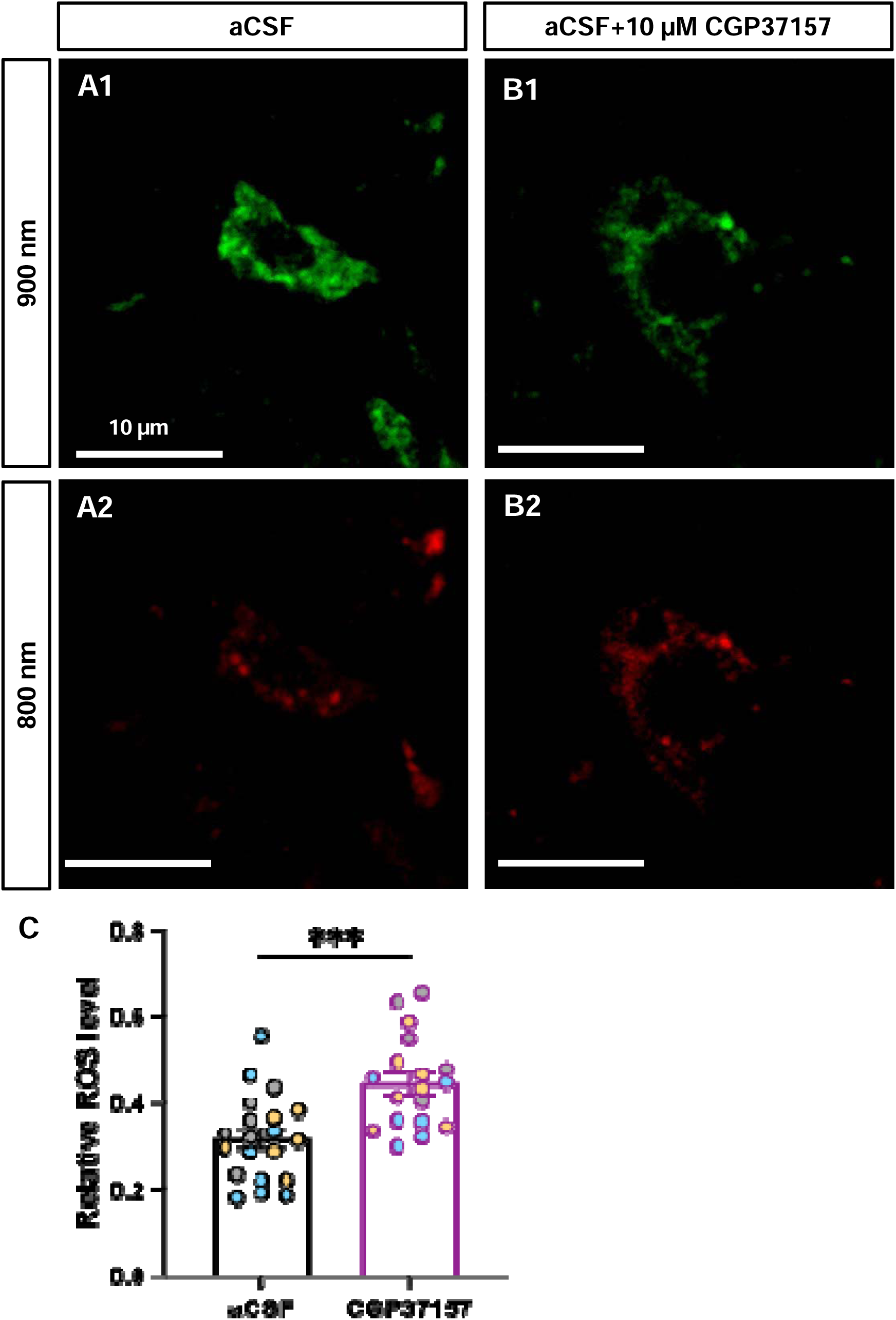
NCLX inhibition-induced mitochondrial Ca^2+^ overload leading to enhanced oxidative levels. (A-B) Representative images of SN dopaminergic neurons expressing roEGFP (4-5-month-old). Neurons were imaged at excitation wavelengths of 900 nm (reduced state, A1 and B1) and 800 nm (oxidized state, A2 and B2) under two conditions: (A) regular aCSF, (B) aCSF supplemented with 10 µM CGP37157. **(C)** Quantification of oxidative stress levels following 60-minute treatment with 10 µM CGP37157. Each data point represents the mean ROS level of an individual neuron, with colors denoting neurons from distinct brain slices. *P*-value was determined using the unpaired t-test. \*\*\**P* < 0.001. Ctrl: n = 20; CGP37157: n = 19, where n represents the number of individual neurons analyzed.

Subsequently, to investigate the impact of elevated ROS levels on mitochondrial trafficking dynamics, we exposed acute brain slice preparations from WT mice (5-7-month-old) to 50 μM menadione (Fig. 6 A-D). Menadione is a widely employed compound known to exacerbate endogenous ROS production in cultured cell systems^27^. Following a 20-minute incubation with 50 µM menadione, the proportion of stationary mitochondria significantly increased from 88.71 ± 1.49% to 96.75 ± 0.86% (*P* = 0.0036, Fig. 6 C). Anterograde moving mitochondria were reduced by nearly 70%, from 6.70 ± 1.14% to 2.20 ± 0.73% (*P* = 0.0202), while retrograde movement also declined (4.59 ± 0.72% (aCSF control), 1.05 ± 0.18% (MED), *P* = 0.0177, Figure 6 A-C, Video S4 and S6). The average velocity of both anterograde and retrograde movement remained unchanged after MED treatment (Fig. 6 D). To rule out the possibility that decreased motility was due to DMSO toxicity, we also evaluated the effect of DMSO treatment on mitochondrial trafficking. The results showed no significant change in the percentage of moving mitochondria and their velocity between aCSF control and DMSO treatment (Fig. 6 C-D, Video S4 and S5). Additionally, we examined the effect of H_2_O_2_ treatment, another reagent commonly used to induce ROS generation and observed similar results on mitochondrial trafficking (Fig. S3). These findings suggest that artificially increasing endogenous ROS levels in WT brain slices can impair mitochondrial motility, particularly affecting anterograde movement, similar to the deficits observed in *Pink1^-/-^* mice.

**Fig. 6.**
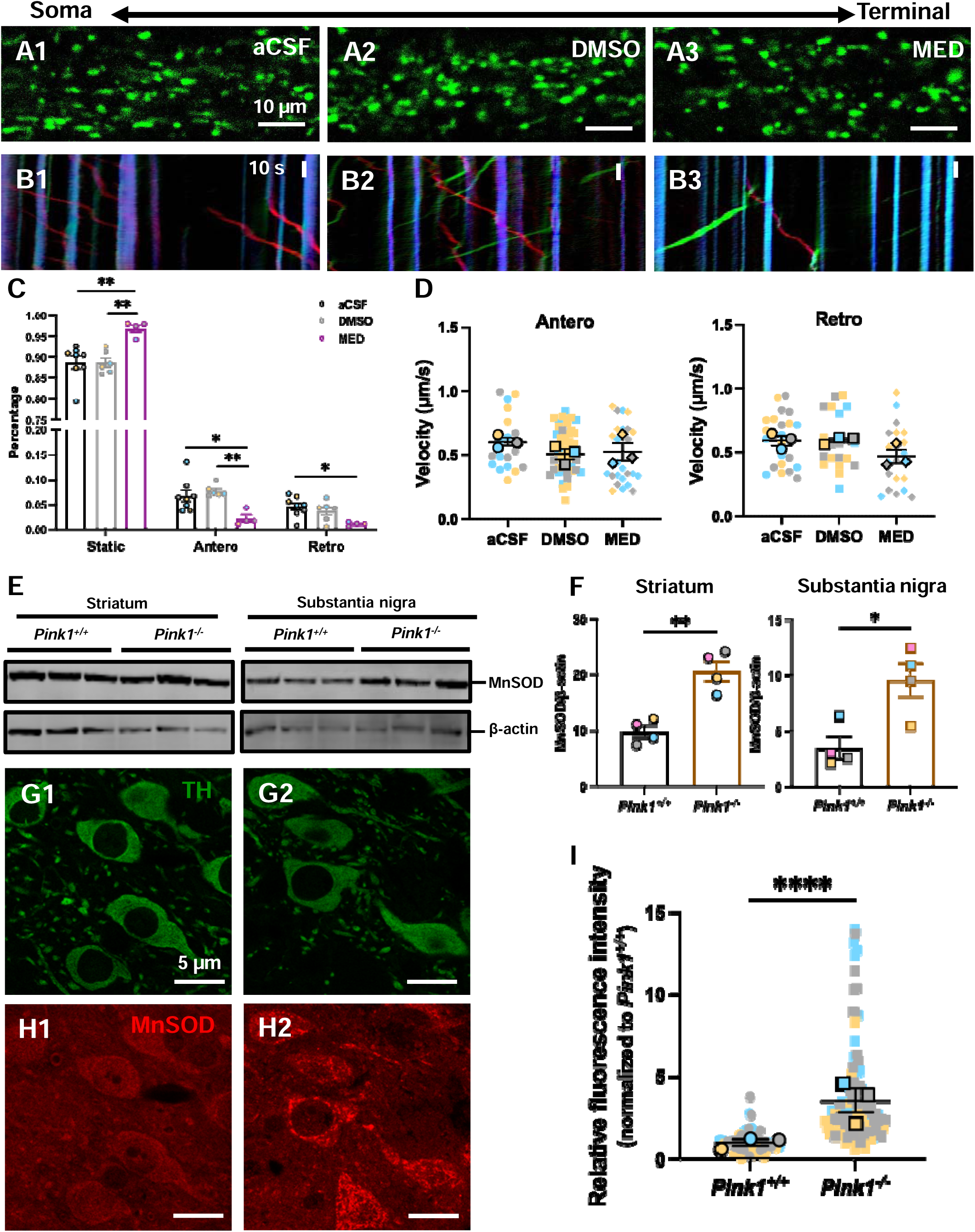
Reduced mitochondrial anterograde movement by ROS elevation. **(A)** Representative images depicting mitochondrial distribution under various treatment conditions at the initial imaging point: (A1) aCSF, (A2) 0.2% DMSO in aCSF, and (A3) 50 μM menadione (MED) in aCSF. **(B)** Kymographs illustrating mitochondrial trafficking patterns following different treatments: (B1) aCSF, (B2) 0.2% DMSO in aCSF, and (B3) 50 μM menadione (MED) in aCSF. **(C)** Quantitative analysis of mitochondrial motility under different treatment conditions (5-7-month-old). *P*-values were determined using the unpaired t-test. \**P* < 0.05, ** *P* < 0.01. Regular aCSF control: N = 3, n = 8; DMSO control: N = 3, n = 6; menadione: N = 3, n = 4. **(D)** Effect of menadione treatment on mitochondrial velocity in anterograde (left) and retrograde (right) directions. **(E)** Representative immunoblots showing MnSOD expression in the striatum (left) and substantia nigra (right) of 5-7-month-old *Pink1*^+/+^ and *Pink1^-/-^* mice. **(F)** Quantification of MnSOD expression levels in striatum (left) and substantia nigra (right) of 5-7-month-old *Pink1*^+/+^ and *Pink1^-/-^* mice. Each color represents an individual animal. *P*-values were determined using the unpaired t-test. \**P* < 0.05, \*\**P* < 0.01. **(G)** Immunofluorescence images of TH-positive dopaminergic neurons in the substantia nigra of 5-7-month-old *Pink1*^+/+^ (G1) and *Pink1^-/-^* (G2) mice. **(H)** Immunofluorescence images of MnSOD in the substantia nigra of 5-7-month-old *Pink1*^+/+^ (H1) and *Pink1^-/-^* (H2) mice. **(I)** Quantification of relative MnSOD fluorescence intensity in dopaminergic neurons of 5-7-month-old *Pink1*^+/+^ and *Pink1^-/-^* mice. Statistical significance was determined using the Mann-Whitney U test. In scatter plots, each data point represents an individual brain slice. In summary plot, each data point represents the mean fluorescence intensity per animal, with colors distinguishing individual mice. \*\*\*\**P* < 0.0001.

Next, we evaluated ROS levels in *Pink1^-/-^* mice. Manganese superoxide dismutase (MnSOD), a crucial mitochondrial antioxidant enzyme, detoxifies oxygen free radicals generated during mitochondrial respiration. Evidence indicates that oxidative stress significantly upregulates MnSOD expression levels^28^. We quantified MnSOD expression levels in the striatum and SNc regions of *Pink1^-/-^* mice and WT littermates. Western blot analysis revealed a significant upregulation of MnSOD levels in 5-7-month-old *Pink1^-/-^* mice compared to WT littermates in both the striatum and SNc regions (Fig. 6 E-F), with an approximate 2-fold increase in striatal MnSOD expression (*P* = 0.0019) and a 1.75-fold elevation in the region of substantia nigra (*P* = 0.0144). Complementary immunostaining corroborated these findings, showing significantly higher MnSOD immunoreactivity in dopaminergic neurons from *Pink1^-/-^* mice compared to WT controls (Fig. 6 H-I). Quantitative analysis revealed an approximate 4-fold increase in relative MnSOD fluorescence intensity in *Pink1^-/-^* mice (*P* < 0.0001). Our previous findings demonstrated decreased mitochondrial coupling efficiency starting at 3-4 months of age and continuing through 14 months^14^. This reduced coupling efficiency directly correlates with increased oxidative stress. Together, these results provide compelling evidence of elevated oxidative levels in PINK1 knockout mice. The excessive ROS generation in acute brain slices from WT mice induced by menadione treatment was associated with impaired mitochondrial motility. These findings underscore a potential pathogenic link between PINK1-mediated mitochondrial dysfunction, dysregulated redox homeostasis, and disrupted mitochondrial trafficking, which may contribute to the degeneration of dopaminergic neurons in PD.

### p38 MAPK pathway hyperactivation by loss of PINK1

We have demonstrated that PINK1 loss causes mitochondrial calcium overload in dopaminergic neurons, resulting in a significant increase in intracellular ROS levels. Oxidative stress plays a pivotal role in dopaminergic neuronal death. Environmental toxins like paraquat and rotenone induce ROS production, which activates the p38 MAPK pathway. This activation not only contributes to mitochondrial dysfunction and apoptosis but also creates a feedback loop that exacerbates oxidative stress, accelerating neurodegeneration. Interventions targeting p38 MAPK have shown promise in disrupting this deleterious cycle^29^. However, the specific mechanisms by which excessive ROS affect mitochondrial movement in neurons remain to be fully elucidated. Previous studies have shown that intracellular ROS can inhibit the function of the motor-adaptor complex that regulates mitochondrial motility through p38 MAPK pathway, thus reducing mitochondrial motility *in vitro*^27^.

We analyzed p38 phosphorylation level in *Pink1^-/-^* mice (6-7-month-old) (Fig. 7 A-B). Immunoblot results showed that p38 phosphorylation levels were approximately threefold higher in *Pink1^-/-^* mice than in WT mice in the striatum (*P* = 0.0286). The substantia nigra region exhibited a similar increase in p38 phosphorylation levels (*P* = 0.0079). Elevated p38 phosphorylation indicates heightened activation of the p38 pathway. Collectively, these results suggest hyperactivation of the p38 MAPK pathway in *Pink1^-/-^* mice.

**Fig. 7.**
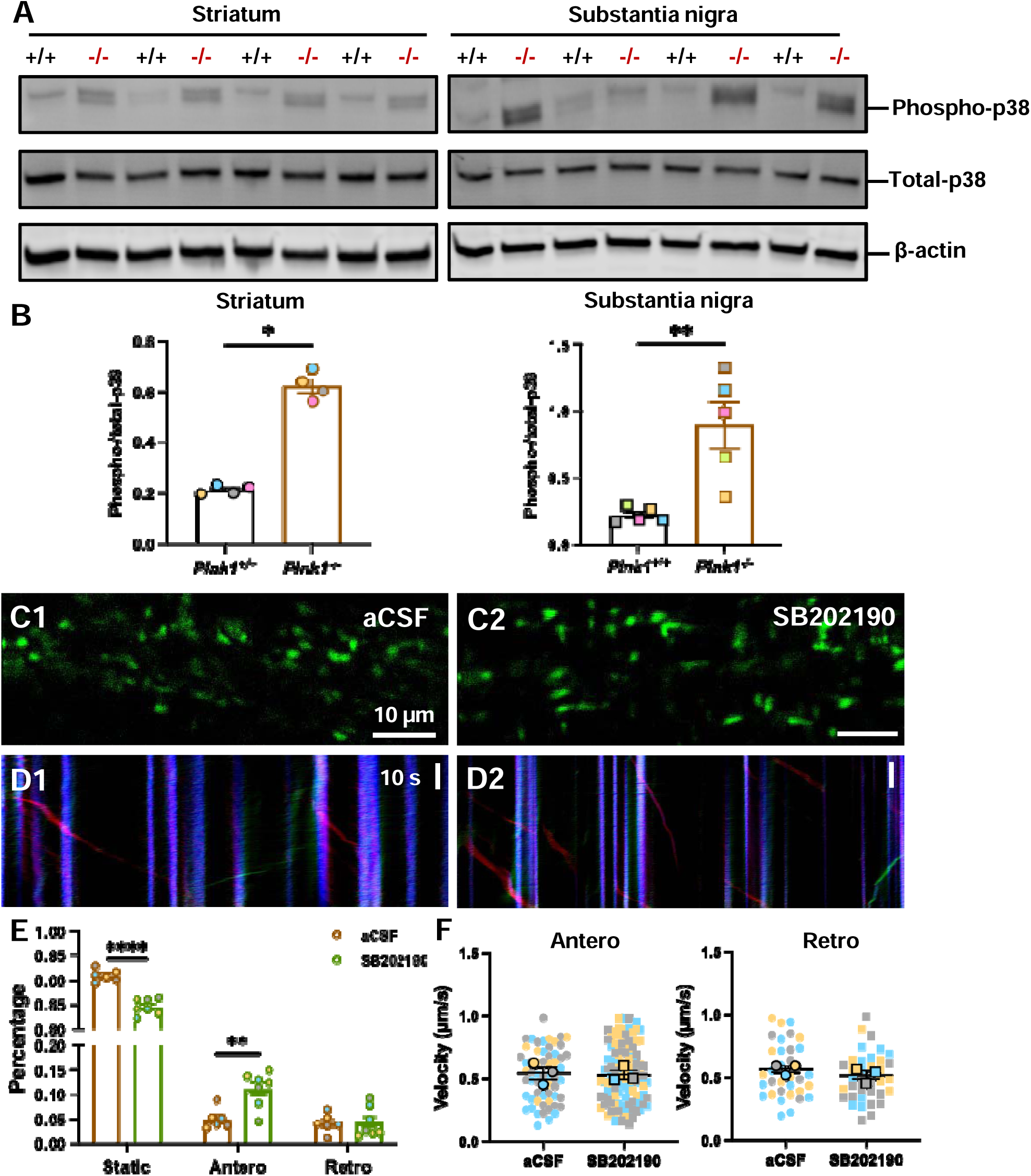
Improved mitochondrial motility in *Pink1^-/-^* mice by inhibiting p38 MAPK hyperactivity. **(A)** Representative immunoblots showing phospho-p38 and total p38 expression levels in the striatum (left) and substantia nigra (right) of 6-7-month-old *Pink1^+/+^* and *Pink1^-/-^* mice. **(B)** Quantification of phospho-p38 and total p38 expression levels in striatum (left) and substantia nigra (right). Each data point represents an individual animal. *P*-values were calculated using Mann-Whitney U-test. \*\**P* < 0.01, \*\*\*\**P* < 0.0001. **(C)** Representative images depicting mitochondrial distribution under different treatment conditions at the initial imaging point: (C1) aCSF, (C2) 50 μM p38 inhibitor (SB202190) in aCSF. **(D)** Kymographs illustrating mitochondrial trafficking patterns: (D1) aCSF, (D2) 50 μM SB202190 treatment. **(E)** Percentage quantification of mitochondrial movement status after SB202190 incubation. *P*-values were determined using the unpaired t-test. \*\**P* < 0.01, \*\*\*\**P* < 0.0001. Regular aCSF control: N = 3, n = 6; SB202190: N = 3, n = 7. **(F)** Impact of SB202190 treatment on mitochondrial velocity in anterograde (left) and retrograde (right) directions. *P*-values were calculated using the Mann-Whitney U-test.

We extended our analysis to examine the phosphorylation levels of two other MAPK family pathways: JNK (Fig. S4) and ERK (Fig. S5). JNK phosphorylation levels were also elevated in *Pink1^-/-^* mice, while ERK phosphorylation levels remained unchanged. Both p38 and JNK pathways play crucial roles in cellular stress responses. The observed hyperactivation of these pathways in *Pink1^-/-^* mice aligns with our previous results and provides additional support for the oxidative stress hypothesis in PINK1 deficiency (Fig. 6 H).

### Improved mitochondrial motility in *Pink1^-/-^* mice by inhibiting p38

To investigate whether inhibiting p38 activity could alleviate mitochondrial trafficking deficits in *Pink1^-/-^* mice, we treated acute brain slices from these mice with the p38 inhibitor SB202190 (Fig. 7 C-F, Video S7-S8). Incubating *Pink1^-/-^* mice brain slices in aCSF containing 50 μM SB202190 for one hour significantly increased mitochondrial motility in dopaminergic neurons (Fig. 7 E). The proportion of immobile mitochondria decreased from 90.93 ± 0.51% to 84.59 ± 0.61% (*P* < 0.0001). The percentage of anterograde-moving mitochondria significantly increased from 4.96 ± 0.86% to 11.12 ± 1.36% (*P* = 0.0037). Retrograde mitochondrial movement was not significantly affected (aCSF: 4.11 ± 0.78% *vs.* SB202190: 4.29 ± 1.15%). Additionally, SB202190 treatment did not alter the velocity of either anterograde or retrograde mitochondrial movement (Fig. 7 F).

We also verified the effectiveness of SB202190 treatment in *Pink1^-/-^* mouse brain slices through immunoblot analysis, which showed a significant ∼62% reduction in p38 phosphorylation (Fig. S6). This substantial decrease in p38 phosphorylation led to a significant rescue of mitochondrial motility defects in dopaminergic neurons of *Pink1^-/-^* mice, particularly by increasing the percentage of anterograde movement. However, SB202190 treatment did not improve the decreased anterograde velocity in *Pink1^-/-^* mice.

We analyzed the protein expression involved in axonal mitochondrial transport (Fig. S7). The adaptor protein Miro1 showed a significant reduction in *Pink1^-/-^* mice compared to WT littermates: approximately a 15% decrease in the striatum (*P* = 0.0286) and approximately a 24% decrease in the substantia nigra (*P* = 0.0317). This reduction in Miro1 may lead to decreased mitochondrial attachment to microtubules and reduced motility. However, we observed no significant changes in the expression of the motor proteins kinesin and dynein in *Pink1^-/-^* mice (Fig. S7).

### Defective mitochondria at dopaminergic neuron terminals in a *Pink1*^-/-^ mice

*Pink1^-/-^* mice exhibited age-dependent deficits in striatal dopamine release and ATP levels^30^. This observation, along with the impaired mitochondrial axonal transport in *Pink1^-/-^* dopaminergic neurons, led us to hypothesize that reduced anterograde movement of mitochondria in *Pink1^-/-^* dopaminergic neurons might result in a depletion of healthy mitochondria at their axon terminals. To test this hypothesis, we employed tyrosine hydroxylase (TH) immunogold electron microscopy to directly assess mitochondrial morphology and quantify mitochondrial numbers. In WT mice (8-9-month-old), dopaminergic neurons exhibited relatively intact mitochondrial cristae, indicating overall healthy mitochondria (Fig. 8 A). In contrast, *Pink1^-/-^* mice displayed notable mitochondrial abnormalities, including rounded mitochondrial shapes, indistinct cristae, and vacuolated internal structures, suggesting mitochondrial fragmentation (Fig. 8 B). In *Pink1^-/-^* mice, there was a significant decrease in the proportion of healthy mitochondria within dopaminergic terminals (*P* = 0.0079). However, the overall percentage of healthy mitochondria in the striatum remained unchanged (Fig. 8 C, *P* = 0.4524). Compared with WT mice, the circularity (*P* < 0.0001) and aspect ratio (*P* < 0.0001) of mitochondria in the dopaminergic neurons of *Pink1^-/-^* mice were significantly reduced (Fig. 8 D), indicating that the mitochondria at the terminals of dopaminergic neurons in *Pink1^-/-^* mice tended to be more rounded and smaller.

**Fig. 8.**
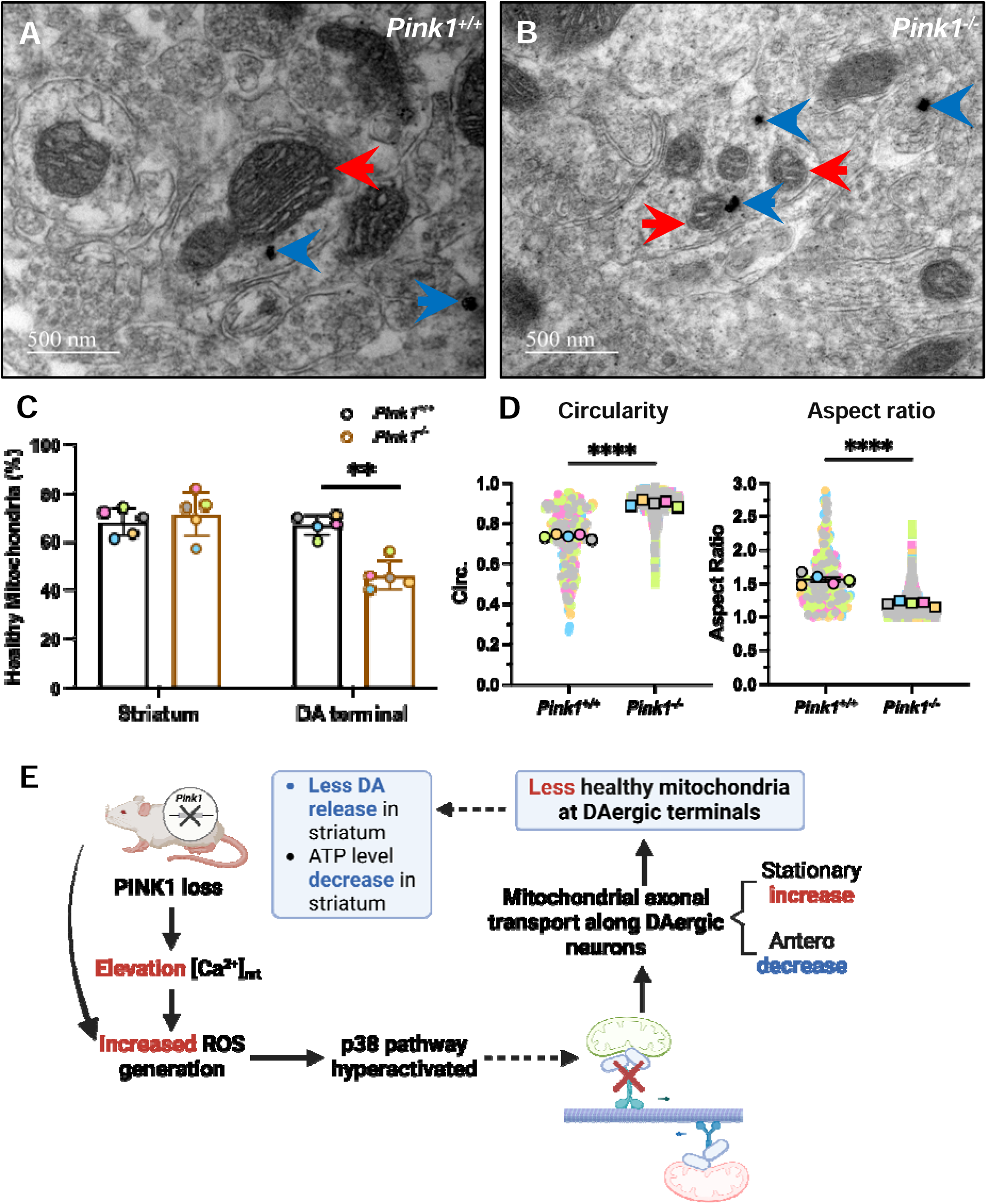
Defective mitochondria at dopamine neuron terminals in *Pink1^-/-^* mice. (A-B) TH immunogold electron micrographs illustrating mitochondrial morphology in dopaminergic terminals of (A) *Pink1^+/+^* and (B) *Pink1^-/-^* mice (8-9-month-old). Blue arrows indicate TH-positive signals in dopamine terminals. Red arrows denote mitochondria within or proximal to dopamine terminals. **(C)** Quantitative analysis of mitochondrial status in dopaminergic terminals and surrounding striatal regions of *Pink1^+/+^* and *Pink1^-/-^*mice. Each data point represents an individual mouse. Statistical significance was determined using the Mann-Whitney U test. \*\**P* < 0.01, N = 5. **(D)** Quantification of mitochondrial circularity and aspect ratio in dopaminergic neuron terminals of *Pink1^+/+^* and *Pink1^-/-^*mice. In scatter plots, each data point represents individual mitochondria. In summary plot, each data point represents the mean value per animal, with colors distinguishing individual mice. *P*-values were calculated using Mann-Whitney U test. \*\*\*\**P* < 0.001, N = 5. Morphometric analysis of mitochondrial circularity and aspect ratio in dopaminergic neuron terminals of *Pink1^+/+^*and *Pink1^-/-^* mice. In scatter plots, each data point represents an individual mitochondrion. In summary plot, each data point represents the mean value per animal, with colors distinguishing individual mice. Statistical analysis was performed using the Mann-Whitney U test. \*\*\*\**P* < 0.001, N = 5. **(E)** Proposed mechanistic model: *Pink1* gene deletion leads to aberrant activation of the p38 pathway, resulting in impaired mitochondrial axonal transport in dopaminergic neurons. Figure was created by BioRender.

Taken together, our findings indicate that the loss of PINK1 leads to a significant increase in mitochondrial calcium levels in the SNc dopaminergic neurons by disrupting specific regulatory mechanisms. This disturbance in mitochondrial calcium homeostasis results in increased oxidative stress and ROS production for p38 signaling cascade activation. Previous studies have shown that p38 can directly phosphorylate kinesin heavy chains (KHCs), thereby inhibiting axonal transport^27,30–33^. Based on these observations, we propose a working model (Fig. 8 G): the increase in ROS caused by loss of PINK1 activates the p38 MAPK pathway, leading to p38-mediated phosphorylation of the anterograde motor protein KIF5. This phosphorylation disrupts the mitochondrial anterograde transport complex, reducing the percentage of mitochondria undergoing anterograde movement. Consequently, the decreased anterograde trafficking of mitochondria results in a lower quantity of healthy mitochondria at the terminals of dopaminergic neurons. This deficit may lead to a reduction in ATP content at the neuron terminals, causing an energetic shortfall and decreased in dopamine release.

## Discussion

Here, we developed and validated a novel approach that synergistically combines Mito-Dendra2 mice, acute brain slice preparations, and 2PLSM to investigate mitochondrial axonal transport within dopaminergic neurons. This strategy allowed us to robustly interrogate mitochondrial transport kinetics in an *ex vivo* setting that more accurately recapitulates the pathophysiological scenario of PD in dopaminergic neurons, owing to the following key advantages: **(i)** Cell-type specificity: We focused specifically on the nigrostriatal dopaminergic neurons, which are the right and most vulnerable neuronal population in PD pathogenesis. **(ii)** Preserved neuronal network: Utilization of acute brain slices from mice provides a more integrated and physiologically relevant neuronal network that closely mimics *in vivo* conditions. Notably, the relative thickness of these slices helps maintain the coherence of the neuronal circuitry, including interactions among neurons, astrocytes, and glial cells, for a considerable duration, despite potential mechanical damage from sectioning. **(iii)** Sustained endogenous neuronal functionality and amenability of pharmacological studies: This approach not only preserves the intrinsic functional activity of dopaminergic neurons for an extended duration but also enables us to perform pharmacological manipulations and assess their effects on neuronal function and mitochondrial dynamics within an integrated *ex vivo* system.

Using this method, we investigated the mitochondrial trafficking alteration in our previous reported *Pink1^-/-^* mice^14^. Our findings reveal that loss of PINK1 causes age-dependent mitochondrial trafficking impairments in dopaminergic neurons, reducing healthy mitochondrial presence at terminals and provides a plausible mechanistic explanation for our previous finding^14^. Corroborating our findings, previous studies have also demonstrated that genetic ablation of *Pink1* in mouse primary cortical and midbrain neurons led to a significant reduction in the proportion of mitochondria undergoing anterograde movement along dendrites^12^. Interestingly, some studies investigating loss of PINK1 in different models or neuronal subtypes have reported contrasting findings. Researches in rat primary hippocampal and *Drosophila* demonstrated that PINK1 deficiency led to an increase in the proportion of motile mitochondria^11,34^. These divergent findings suggest that the effects of PINK1 deficiency on mitochondrial dynamics are context-dependent, varying across species, cell types, and developmental stages. It is in this context that our approach to study mitochondria in dopaminergic neurons in their endogenous architecture is more specific and relevant to PD.

Regarding the underlying mechanism of these deficits, we observed elevated ROS levels and p38 hyperactivation in *Pink1^-/-^* mice. Furthermore, we demonstrated that pharmacological inhibition of p38 rescued mitochondrial motility. Interestingly, our results are consistent with previous studies that suggested a potential mechanism of how p38 activation specifically impacts mitochondrial axonal transport dynamics: without affecting kinesin protein level, enhanced p38 activity can lead to an increase in the phosphorylation of the KIF5 protein, a key component of the kinesin motor complex, which results in a reduced binding affinity to microtubules^27,30–33^.

Collectively, our studies introduce a novel approach to specifically investigate mitochondrial trafficking dynamics within dopaminergic neurons in Parkinson’s disease mouse models. Applying this approach to *Pink1^-/-^* mice, we observed a unique phenotype of mitochondrial trafficking in dopaminergic neurons that differed from previous findings. This method can be further refined and adapted for other disease contexts and animal models in future studies. By integrating this approach with various tool mouse models, it offers the potential to investigate the transport dynamics of diverse organelles or proteins beyond mitochondria in different cellular environments.

## Methods

### Animals

Mice were housed in a temperature and humidity controlled specific pathogen-free (SPF) facility under a 12-hour light/dark cycle schedule with food and water available ad libitum. Mice were on FVB/NTac mouse background (Taconic). All experimental protocols were approved by the Animal Care and Use Committees of Thomas Jefferson University, Peking University, and The University of Georgia.

### Construction, modification and confirmation of TH-MitoDendra2-BAC

BAC clone covering the entire tyrosine hydroxylase (*Th*) gene was obtained from GENSAT RPCI23-350E13 (http://genome.ucsc.edu/cgi-bin/hgTracks?clade=mammal&org=Mouse&db=mm9&position=RP23-350E13).

Dendra2-SV40 polyA from Clontech pCMV vector was ligated with a mitochondrial targeting sequence^15^. The mitochondrial targeting sequence (MTS) was from human subunit 8A of cytochrome c oxidase (COX8A): 5’ATGTCCGTCCTGACGCCGCTGCTGCTGCGGGGCTTGACAGGCTCGGCCCGGCGGCTCCCAGTGCCGCGCGCCAAGATCCATTCGTTGGGGGATCCACCGGTCGCCACC3’. Mito-Dentra2 was inserted to the exon1 of *Th* gene with homologous recombination. Homologous arms, A arm (642 bp) and B arm (538 bp), from the first exon of *Th* were ligated to 5’ and 3’ ends of the Mito-Dendra2 cassette by PCR amplification respectively and inserted into the shuttle vector following the standard protocol^16^. The shuttle vectors were introduced into the RP23-350E13 competent cells to construct the co-integrated BACs using electroporation. Recombinants were identified by chloramphenicol and ampicilin resistant selection followed by PCR analysis^16^. Purified TH-MitoDendra2 BACs were used to generate TH-MitoDendra2 transgenic mice.

### TH-MitoDendra2-BAC transgenic mice

TH-MitoDendra2-BACs were used for pronuclear injection in the Taconic FVB/NTac background. Founders and progeny carrying the TH-MitoDendra2 transgene were identified by PCR analysis of tail DNA and direct sequencing. The expression pattern of Mito-Dendra2 was confirmed using TH immunofluorescence in TH-MitoDendra2-BAC transgenic mice. The transgenic mice were maintained by breeding with wild-type (WT) Taconic FVB/NTac mice.

### Immunofluorescence

Mice were anesthetized with ketamine/xylazine (100 mg/10 mg per kg) and subsequently perfused with ice-cold 4% paraformaldehyde in phosphate-buffered saline (PBS) and postfixed for 24 hours. Coronal sections, 30 μm thick, were obtained using a Thermo Scientific Microm HM550 Cryostat. Serial sections were collected into 10 wells, with each well containing every 10th section. Floating sections were stored in antifreeze buffer (30% glycerol and 30% ethylene glycol in PBS) at −20°C. The sections were stained with a polyclonal anti-tyrosine hydroxylase (TH) antibody (Millipore, MAB5280) at a 1:1000 dilution for 36 hours at 4°C, followed by incubation with a Dylight 594-conjugated secondary antibody (1:500, Life Technologies). Fluorescent images were captured using a Leica SP3800 laser scanning confocal microscope.

### Acute sagittal slice preparation

Animals were anesthetized with either isoflurane (1-2%) or ketamine/xylazine (100 mg/10 mg per kg) and underwent transcardiac perfusion with ice-cold, pre-oxygenated cutting solution containing (in mM): 125 NaCl, 2.5 KCl, 26 NaHCO3, 3.7 MgSO4, 0.3 KH2PO4, and 10 glucose, pH 7.4). The brains were immediately dissected out in cold, pre-oxygenated cutting solution. Sagittal nigrostriatal projection slices were sectioned at a thickness of 300 μm using a vibrating microtome (VT1200, Leica, Solms, Germany) with an angle of approximately 21°. The slices were recovered in oxygenated artificial cerebrospinal fluid (aCSF) containing (in mM): 125 NaCl, 2.5 KCl, 26 NaHCO_3_, 2.4 CaCl_2_, 1.3 MgSO_4_, 0.3 KH_2_PO_4_, 10 glucose, 2 HEPES, pH7.4, at room temperature for > 30 minutes^14^.

### Two-photon laser microscopy imaging of mitochondrial trafficking in brain slices

Parasagittal slices containing nigrostriatal projections were obtained from TH-MitoDendra2 transgenic mice. The brain slices were positioned within a recording chamber and superfused with oxygenated aCSF at a flow rate of 1.5 mL/min. Optical imaging of Dendra signals were acquired using an Ultima multiphoton laser scanner (Bruker) for BX51/61 Olympus microscope with a Titanium-sapphire laser (Chameleon-Ultra2, Coherent Laser Group), equipped with a 20× 1.0 NA water immersion objective (XLUMPLFL20XW, Olympus). The Dendra2 protein was excited at a wavelength of 900 nm, and its fluorescence was collected at 500-560 nm. Images were captured in 8-bit resolution and 4 μs dwell time, with regions of interest set at 512 × 512 pixels. The sampling rate ranged from 0.2 to 0.5 seconds per frame. All experiments were conducted at 33-35 °C.

Images were processed using the open-source software Fiji (NIH) and MATLAB (Version 8.5.0 R2015a, MathWorks). Intensity-based alignment was performed to register images at different time points. To measure mitochondrial motility velocity, the plugins KymographClear and KymographDirect were used to generate kymographs representing moving distance versus moving duration. The number and percentage of mitochondria in different states—static, retrograde, and anterograde—were quantified and measured. For each experiment, multiple kymographs were generated from the same field in order to capture most moving mitochondria. Overall velocity was determined by calculating the ratio of total displacement to observation duration. For mitochondria exhibiting bidirectional movement, comprehensive trajectory analysis was performed using kymographs, where directional changes were identified by slope inversions, enabling velocity calculations for individual directional segments.

### Stereotaxic injection of AAV

3-4-month-old male mice of *Pink1*^+/+^and *Pink1*^-/-^ were anesthetized with isoflurane (1-2%) or ketamine/xylazine (100 mg/10 mg/kg). A small craniotomy, approximately 0.8 mm in diameter, was made over the right midbrain (−3.4 mm caudal, +1.0 mm lateral). To achieve expression of mito-GCaMP6 in dopaminergic neurons, 0.1-0.2 µl of AAV-GCaMP6-mito virus (10^13 vg/ml, a gift from Dr. James Surmeier’s lab at Northwestern University) was pressure-injected through a syringe into the midbrain at a depth of -4.2 mm ventral from the dura surface. Following the injections, the incisional wound on the skull was sealed with tissue adhesive (3M Vetbond). The mice were placed on a heating pad until fully recovered and returned to the isolation room. Carprofen (5 mg/kg, Sigma Aldrich) was administered intraperitoneally once per day for at least 3 days to reduce pain.

### 2PLSM mito-GCaMP6 imaging in acute brain slices

Mitochondrial Ca^2+^ levels were measured with the mitochondrially targeted Ca^2+^-sensitive probe mito-GCaMP6^17^. Two-three weeks after AAV virus injection, mice were prepared for coronal brain slice sectioning following transcardiac perfusion with ice-cold, oxygenated cutting solution. After a recovery period of 30-45 minutes, the coronal brain slices were placed in a chamber perfused with oxygenated aCSF. To measure free Ca^2+^ levels in mitochondria, the slices were sequentially treated with the following solutions: (1) regular aCSF, (2) 500 µM EGTA + aCSF without Ca^2+^ for 30-40 minutes, (3) 500 µM EGTA + 1 µM ionomycin + aCSF without Ca^2+^ for 30 minutes, and (4) 1 µM Ionomycin + aCSF with high Ca^2+^ (3 mM) for 20 minutes. Time series images were acquired every 5 minutes to determine the minimal and maximal fluorescence intensity. The fluorescence intensity at the end of treatment (2) was defined as the minimum fluorescence (Fmin), and the fluorescence intensity at the end of treatment (4) was defined as the maximum fluorescence (Fmax). The relative free calcium level was calculated using the equation: (F - Fmin) / (Fmax - Fmin).

### Immunoblot

Tissues were homogenized and lysed with RIPA buffer (Millipore Sigma, R0278) supplemented with fresh protease inhibitors (Roche, 11873580001) and phosphatase inhibitors (Roche, 04906845001). The homogenate was further lysed for 1 hour at 4°C and cleared by centrifugation at 12,000 × g for 15 minutes at 4°C. The supernatants were collected, and protein concentrations were determined using a BCA assay kit (Thermo Scientific Pierce, 23227). Protein samples (30-50 μg) were separated using Mini-PROTEAN® TGX™ Precast Gels (Bio-Rad). After blocking, the blots were incubated with primary antibodies overnight at 4°C. Following washing with 1× TBST (0.1% Tween-20), the blots were incubated with secondary antibodies (1:10000, IRDye® 800RD, Li-COR 926-32211; IRDye® 680RD, Li-COR, 926-68070) for 1 hour and visualized using the Odyssey Infrared Imager (Li-COR, 9120). The relative protein abundance was quantified using Image Studio software (Li-COR).

### TH-Immunogold labeling and electron microscopy

Mice were perfused intracardially with freshly prepared fixatives. The fixative consisted of 3.75% acrolein and 2% paraformaldehyde (PFA) in 0.1 M phosphate buffer, pH 7.4 (PB). After fixation, brains were sectioned at 50 µm in PB using a vibratome subsequently fixed at 4°C overnight in 4% PFA/PB. Sections were treated with 50 mM glycine in 1× PBS for 30 minutes to reduce free aldehydes and rinsed extensively with PB three times. The sections were subsequently permeabilized with Tris-buffered saline (TBS) containing 3% goat serum, 1% bovine serum albumin (BSA), and 0.04% Triton X-100 for 30 minutes. The primary mouse anti-TH antibody (#MAB5280, Millipore Sigma) was diluted 1:100 in PBS with 3% goat serum and 1% BSA. The sections were incubated with the primary antibody at 4°C for 48 hours. After washing three times (10 minutes each) with PBS, the sections were labeled with 0.8 nm gold-conjugated goat anti-mouse secondary antibody (#800.177, Aurion) diluted in a buffer containing 3% goat serum, 0.8% BSA, and 0.1% cold fish gelatin in PBS. The sections were incubated with the secondary antibody at 4°C for 48 hours. Following one wash with dilution buffer and six washes with PBS, sections were post-fixed with 2.5% glutaraldehyde in PBS. After several washes with PBS, the sections were exposed to Enhancement Conditioning Solution (ECS; #500.055, Aurion) diluted 1:10 with ddH2O, for four times at 10 minutes each. Ultra-small gold particles were enlarged using R-Gent SE-EM Silver Enhancement Reagents (#500.044, Aurion) for 150 minutes. The sections were rinsed in ECS before being transferred to PBS. Tissue sections were transferred to PBS in Coors dishes and fixed in 1% osmium tetroxide in PB for 60 minutes on ice. After rinsing in ddH2O, the sections were sequentially dehydrated in 30%, 50%, 70%, 85%, and 95% ethanol solutions, followed by two exposures to 100% ethanol and two exposures to acetone. Tissue sections were incubated overnight in a 1:1 mixture of epoxy resin (#14120, EMbed 812, Electron Microscopy Sciences) and acetone before being transferred to pure EMbed 812 for 3 hours (three times, 1 hour each). The sections were embedded in EMbed 812 and polymerized for 24 hours at 65°C. Plastic blocks were trimmed and sectioned with a Leica ultramicrotome (EM UC7, Leica) equipped with a 35° diamond knife (Diatome). Serial ultrathin sections (75 nm) were collected on single-slot formvar-coated copper grids. After post-staining with uranyl acetate and lead citrate, the grids were examined under a transmission electron microscope (Tecnai G^2^ Spirit BioTWIN, FEI) equipped with a digital camera (Orius 832, Gatan) at 120kV.

### Antibodies and Reagents

The antibodies and reagents used in the study include: Anti-Tyrosine Hydroxylase (rabbit, 1:5000 for IF, Pel-Freez, P40101-0), Anti-Tyrosine Hydroxylase (mouse, 1:500 for IF, Millipore, MAB5280), Anti-Kinesin heavy chain (mouse, 1:500 for WB, Millipore, MAB1614), Anti-Dynein medium chain (mouse, 1:1000 for WB, Millipore, MAB1618), Anti-Miro1 (rabbit, 1:1000 for WB, Antibodies_online, ABIN635090), MAPK Family Antibody (rabbit, 1:1000 for WB, Cell signaling, 9926), Phospho-MAPK Family Antibody (rabbit, 1:1000 for WB, Cell signaling, 9910), Anti-MnSOD (rabbit, 1:1000 for WB, Abcam, ab68155), Anti-MnSOD (rabbit, 1:100 for IF, Abcam, ab13534), and Anti-β-actin (rabbit, 1:1000 for WB, Cell signaling, 4970T). Secondary antibodies used were DyLight 594 Antibody (Goat anti-rabbit, 1:500 for IF, Invitrogen, 35560), Alexa Fluor 488nm (Goat anti-mouse, 1:1000 for IF, Invitrogen, A-11029), Alexa Fluor 594nm (Goat anti-rabbit, 1:1000 for IF, Invitrogen, A-11012), IRDye® 800CW (Goat anti-rabbit, 1:5000 for WB, Li-COR, 926-32211), and IRDye® 680RD (Goat anti-mouse, 1:5000 for WB, Li-COR, 926-68070). Additional reagents included A23187 (Sigma-Aldrich, C7522), CGP37157 (Sigma-Aldrich, C8874), menadione (Sigma-Aldrich, M5625), and ionomycin (Sigma-Aldrich, I3909).

### Data and Statistical Analysis

Data were presented as mean ± SEM unless otherwise specified. Statistical analyses were conducted using Prism 9.0 (GraphPad), including the unpaired t test, Mann–Whitney test and one-way ANOVA. Differences were considered statistically significant when *P* < 0.05. Statistical analysis was based on the number of slices or experiments unless otherwise noted. “N” denotes the number of mice, and “n” denotes the number of experiments.

## Data availability

Data generated or analyzed during this study are all included in the published article and its Supplymentary files. All other data are available from the corresponding authors upon reasonable request.

## Acknowledgements

The authors wish to thank Dr. Ying-Chun Hu, Zhen-Yang Kong, and Hong-Zhang Zhou for their expert technical assistance with EM sample preparation and image analysis at the Core Facilities of the School of Life Sciences, Peking University, Dr. D. James Surmeier at Northwestern University for providing the AAV-TH-GCamp6 virus and TH-Mito-roGFP mice, Dr. Jerry Cheng (New York Institute of Technology) for his valuable suggestions on the statistical analysis, and Dr. Shiquan Cui at UGA for his feedback on the manuscript.

## Author Contributions

Jingyu Zhao, Conceptualization, Data curation, Formal analysis, Validation, Visualization, Writing – original draft, Writing – review and editing; Yuanxin Chen, Conceptualization, Data curation, Formal analysis, Validation; Lianteng Zhi, Qing Xu, Data curation; Hui Zhang, Chenjian Li, Conceptualization, Data curation, Formal analysis, Funding acquisition, Investigation, Methodology, Project administration, Supervision, Validation, Writing – review and editing.

## References

1. Poewe, W. et al. Parkinson disease. Nat Rev Dis Primers 3, 1–21 (2017).

2. Malpartida, A. B., Williamson, M., Narendra, D. P., Wade-Martins, R. & Ryan, B. J. Mitochondrial Dysfunction and Mitophagy in Parkinson’s Disease: From Mechanism to Therapy. Trends Biochem Sci 46, 329–343 (2021).

3. Winklhofer, K. F. & Haass, C. Mitochondrial dysfunction in Parkinson’s disease. Biochim Biophys Acta 1802, 29–44 (2010).

4. Pryde, K. R., Smith, H. L., Chau, K.-Y. & Schapira, A. H. V. PINK1 disables the anti-fission machinery to segregate damaged mitochondria for mitophagy. J Cell Biol 213, 163–171 (2016).

5. Exner, N., Lutz, A. K., Haass, C. & Winklhofer, K. F. Mitochondrial dysfunction in Parkinson’s disease: molecular mechanisms and pathophysiological consequences. EMBO J 31, 3038–3062 (2012).

6. Vilain, S. et al. The yeast complex I equivalent NADH dehydrogenase rescues pink1 mutants. PLoS Genet 8, e1002456 (2012).

7. Park, J.-S., Davis, R. L. & Sue, C. M. Mitochondrial Dysfunction in Parkinson’s Disease: New Mechanistic Insights and Therapeutic Perspectives. Curr Neurol Neurosci Rep 18, 21 (2018).

8. Kostic, M. et al. PKA Phosphorylation of NCLX Reverses Mitochondrial Calcium Overload and Depolarization, Promoting Survival of PINK1-Deficient Dopaminergic Neurons. Cell Rep 13, 376–386 (2015).

9. Weihofen, A., Thomas, K. J., Ostaszewski, B. L., Cookson, M. R. & Selkoe, D. J. Pink1 forms a multiprotein complex with miro and milton, linking Pink1 function to mitochondrial trafficking. Biochemistry 48, 2045–2052 (2009).

10. Zhu, X.-H. et al. Quantitative imaging of energy expenditure in human brain. Neuroimage 60, 2107–2117 (2012).

11. Wang, X. et al. PINK1 and Parkin target miro for phosphorylation and degradation to arrest mitochondrial motility. Cell 147, 893–906 (2011).

12. Das Banerjee, T., et al. PINK1 regulates mitochondrial trafficking in dendrites of cortical neurons through mitochondrial PKA. J Neurochem 142, 545–559 (2017).

13. Sheng, Z. H. Mitochondrial trafficking and anchoring in neurons: New insight and implications. Journal of Cell Biology vol. 204 1087–1098 Preprint at 10.1083/jcb.201312123 (2014).

14. Zhi, L. et al. Loss of PINK1 causes age-dependent decrease of dopamine release and mitochondrial dysfunction. Neurobiol Aging 75, 1–10 (2019).

15. Wang, X. et al. Amyloid-beta overproduction causes abnormal mitochondrial dynamics via differential modulation of mitochondrial fission/fusion proteins. Proc Natl Acad Sci U S A 105, 19318–19323 (2008).

16. Gong, S., Yang, X. W., Li, C. & Heintz, N. Highly efficient modification of bacterial artificial chromosomes (BACs) using novel shuttle vectors containing the R6Kgamma origin of replication. Genome Res 12, 1992–1998 (2002).

17. Zampese, E. et al. Ca2+ channels couple spiking to mitochondrial metabolism in substantia nigra dopaminergic neurons. Science Advances 8, 8701 (2022).

18. Mangeol, P., Prevo, B. & Peterman, E. J. G. KymographClear and KymographDirect: two tools for the automated quantitative analysis of molecular and cellular dynamics using kymographs. Mol Biol Cell 27, 1948–1957 (2016).

19. Schwarz, T. L. Mitochondrial trafficking in neurons. Cold Spring Harb Perspect Biol 5, a011304 (2013).

20. Liu, W. et al. Pink1 regulates the oxidative phosphorylation machinery via mitochondrial fission. Proc Natl Acad Sci U S A 108, 12920–12924 (2011).

21. Wang, X. & Schwarz, T. L. The mechanism of Ca2+-dependent regulation of kinesin-mediated mitochondrial motility. Cell 136, 163–174 (2009).

22. Giorgi, C., Marchi, S. & Pinton, P. The machineries, regulation and cellular functions of mitochondrial calcium. Nat Rev Mol Cell Biol 19, 713–730 (2018).

23. Ruiz, A., Alberdi, E. & Matute, C. CGP37157, an inhibitor of the mitochondrial Na+/Ca2+ exchanger, protects neurons from excitotoxicity by blocking voltage-gated Ca2+ channels. Cell Death Dis 5, e1156 (2014).

24. Santulli, G., Xie, W., Reiken, S. R. & Marks, A. R. Mitochondrial calcium overload is a key determinant in heart failure. Proc Natl Acad Sci U S A 112, 11389–11394 (2015).

25. Peng, T.-I. & Jou, M.-J. Oxidative stress caused by mitochondrial calcium overload. Ann N Y Acad Sci 1201, 183–188 (2010).

26. Msackyi, M., Chen, Y., Tsering, W., Wang, N. & Zhang, H. Dopamine Release Neuroenergetics in Mouse Striatal Slices. Int J Mol Sci 25, 4580 (2024).

27. Debattisti, V., Gerencser, A. A., Saotome, M., Das, S. & Hajnóczky, G. ROS Control Mitochondrial Motility through p38 and the Motor Adaptor Miro/Trak. Cell Reports 21, 1667–1680 (2017).

28. Candas, D. & Li, J. J. MnSOD in oxidative stress response-potential regulation via mitochondrial protein influx. Antioxid Redox Signal 20, 1599–1617 (2014).

29. He, J., Zhong, W., Zhang, M., Zhang, R. & Hu, W. P38 Mitogen-activated Protein Kinase and Parkinson’s Disease. Translational Neuroscience 9, 147 (2018).

30. Padzik, A. et al. KIF5C S176 Phosphorylation Regulates Microtubule Binding and Transport Efficiency in Mammalian Neurons. Frontiers in cellular neuroscience 10, (2016).

31. Brady, S. T. & Morfini, G. A. Regulation of motor proteins, axonal transport deficits and adult-onset neurodegenerative diseases. Neurobiology of disease 105, 273–282 (2017).

32. Morfini, G. A. et al. Inhibition of Fast Axonal Transport by Pathogenic SOD1 Involves Activation of p38 MAP Kinase. PLOS ONE 8, e65235 (2013).

33. Berth, S. H. & Lloyd, T. E. Disruption of axonal transport in neurodegeneration. The Journal of Clinical Investigation 133, (2023).

34. Liu, S. et al. Parkinson’s disease-associated kinase PINK1 regulates Miro protein level and axonal transport of mitochondria. PLoS Genet 8, e1002537 (2012).

