## Supplementary material for "Loss of PINK1 Causes Age-dependent Mitochondrial Trafficking Deficits in Nigrostriatal Dopaminergic Neurons Through Aberrant Activation of p38 MAPK": Fig. S

### Supplemental files

Zhao J *et al.* Fig. S1

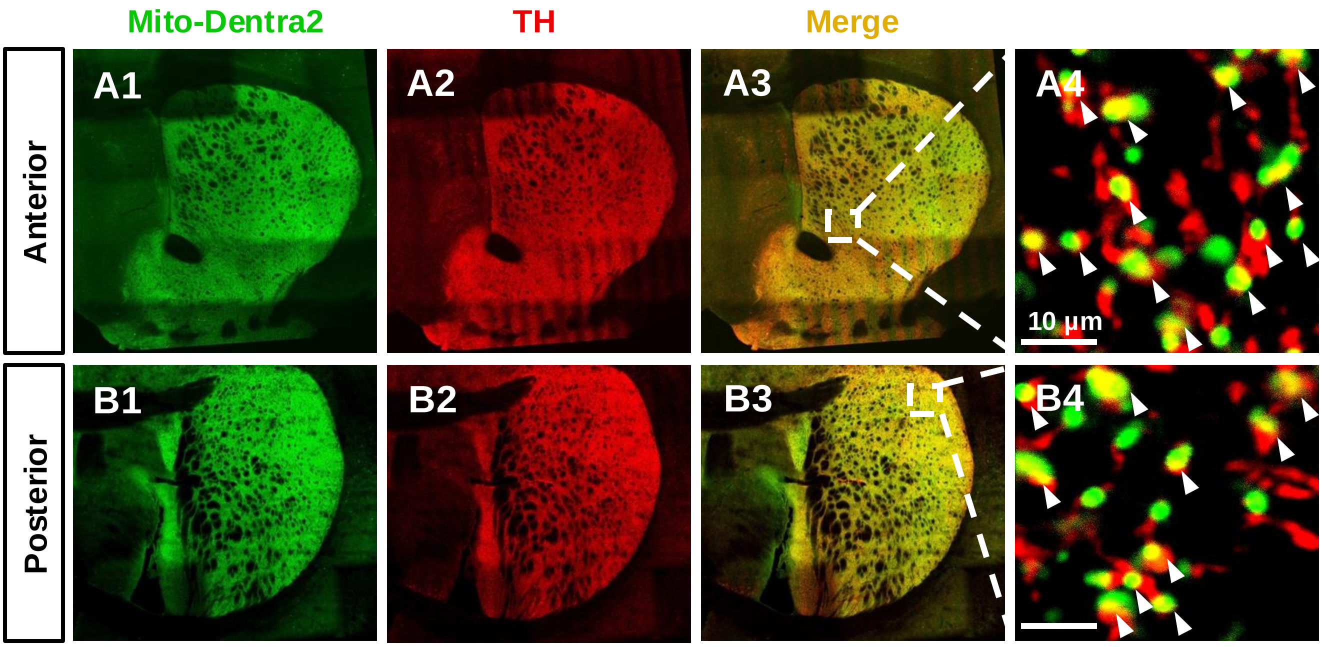


**Fig. S1 Mito-Dendra2 expression in dopaminergic neuron terminals within the striatum of TH-Mito-Dendra2 (MD) BAC transgenic mice. (A)** Mito-Dendra2 expression pattern in the anterior striatum: A1-A3: Low-magnification micrographs demonstrating co-localization of tyrosine hydroxylase (TH, red) and Mito-Dendra2 (green) signals. A4: High-magnification image showing Mito-Dendra2 fluorescence (green) within TH-positive terminals (red) in the striatum. **(B)** Mito-Dendra2 expression pattern in the posterior striatum: B1-B3: Low-magnification micrographs demonstrating co-localization of tyrosine hydroxylase (TH, red) and Mito-Dendra2 (green) signals. B4: High-magnification image showing Mito-Dendra2 fluorescence (green) within TH-positive terminals (red) in the striatum.

Zhao J *et al.* Fig. S2

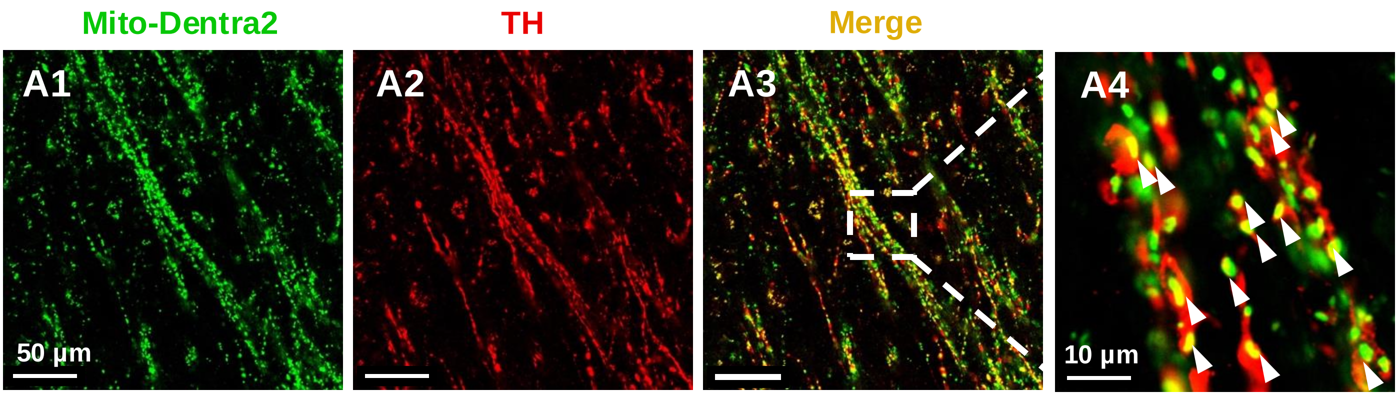


**Fig. S2 Visualization of continuous nigrostriatal axonal projections using Mito-Dendra2 in TH-Mito-Dendra2 transgenic mice. (A)** Representative images depicting Mito-Dendra2 fluorescence in distinct axonal projection bundles.

**Zhao J *et al.* Fig. S3**

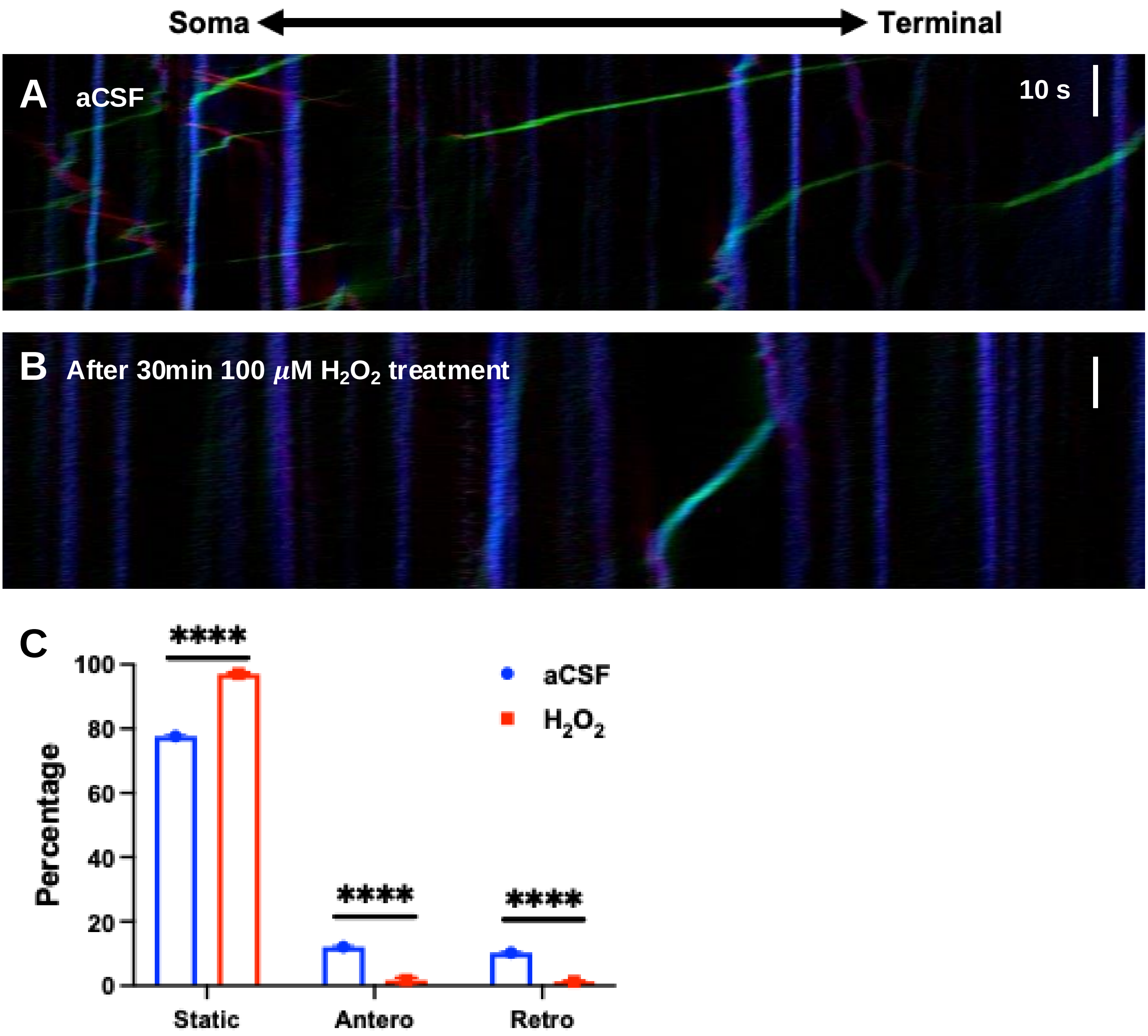


**Fig. S3** Representative kymographs illustrating mitochondrial trafficking patterns before **(A)** and after 100 $\mu$M $H_2O_2$ treatment (**B**). (**C**) Quantification of mitochondrial motility states after $H_2O_2$ treatment (1-month-old mice). P-values were determined using the Mann-Whitney U test.

Zhao J *et al.* Fig. S4

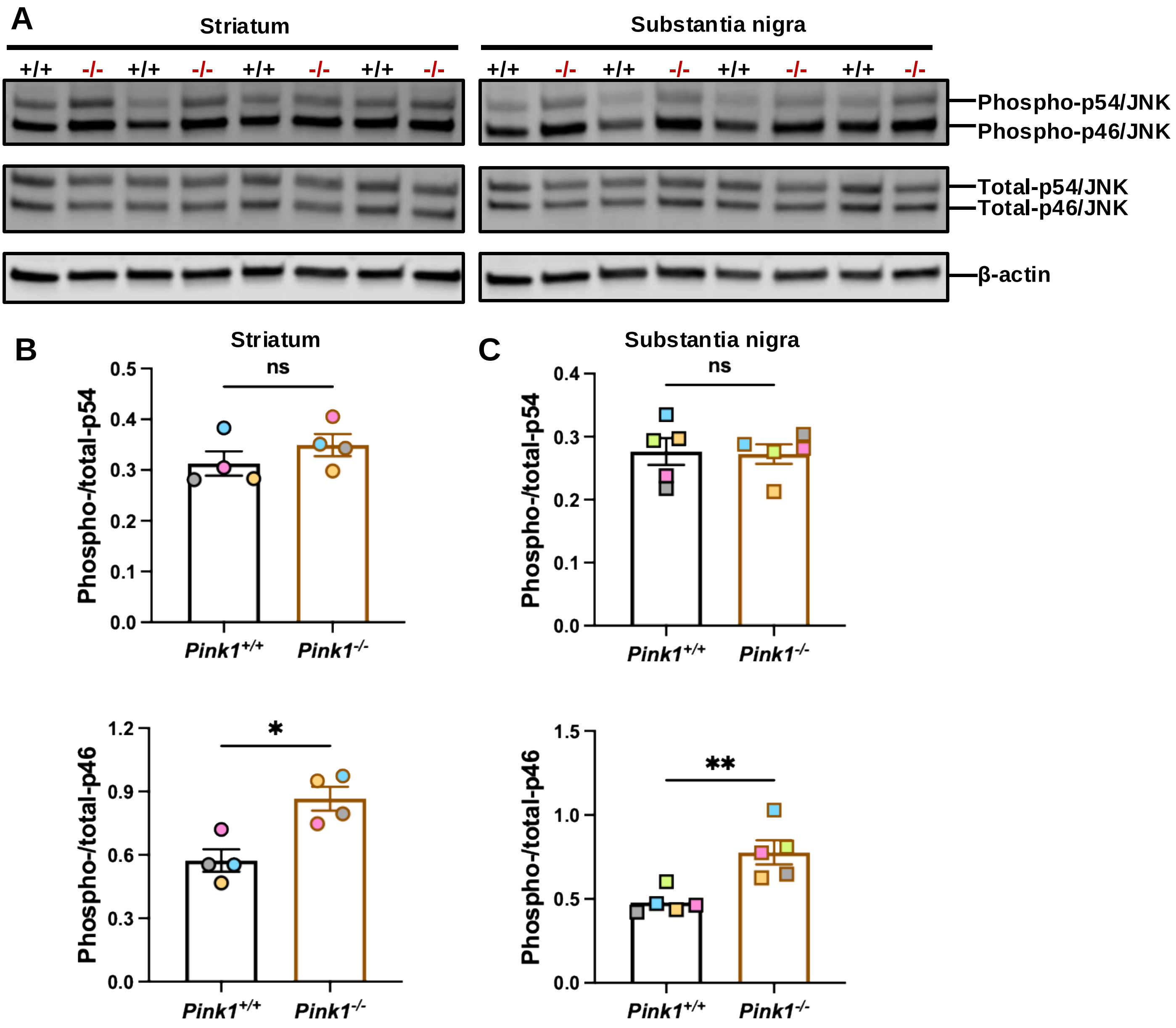


**Fig. S4 Hyperactivation of JNK pathways in *Pink1*$^{-/-}$ mice. (A)** Representative immunoblots showing phospho-JNK and total JNK expression levels in the striatum (left) and substantia nigra (right) of 5-7-month-old *Pink1*$^{+/+}$ and *Pink1*$^{-/-}$ mice. **(B)** Quantification of phospho-JNK and total JNK expression in the striatum. Statistical significance was determined using the Mann-Whitney U test. $^{*}P < 0.05$. **(C)** Quantification of phospho-JNK and total JNK expression in the substantia nigra. Statistical significance was determined using the Mann-Whitney U test. $^{**}P < 0.01$.

Zhao J *et al.* Fig. S5

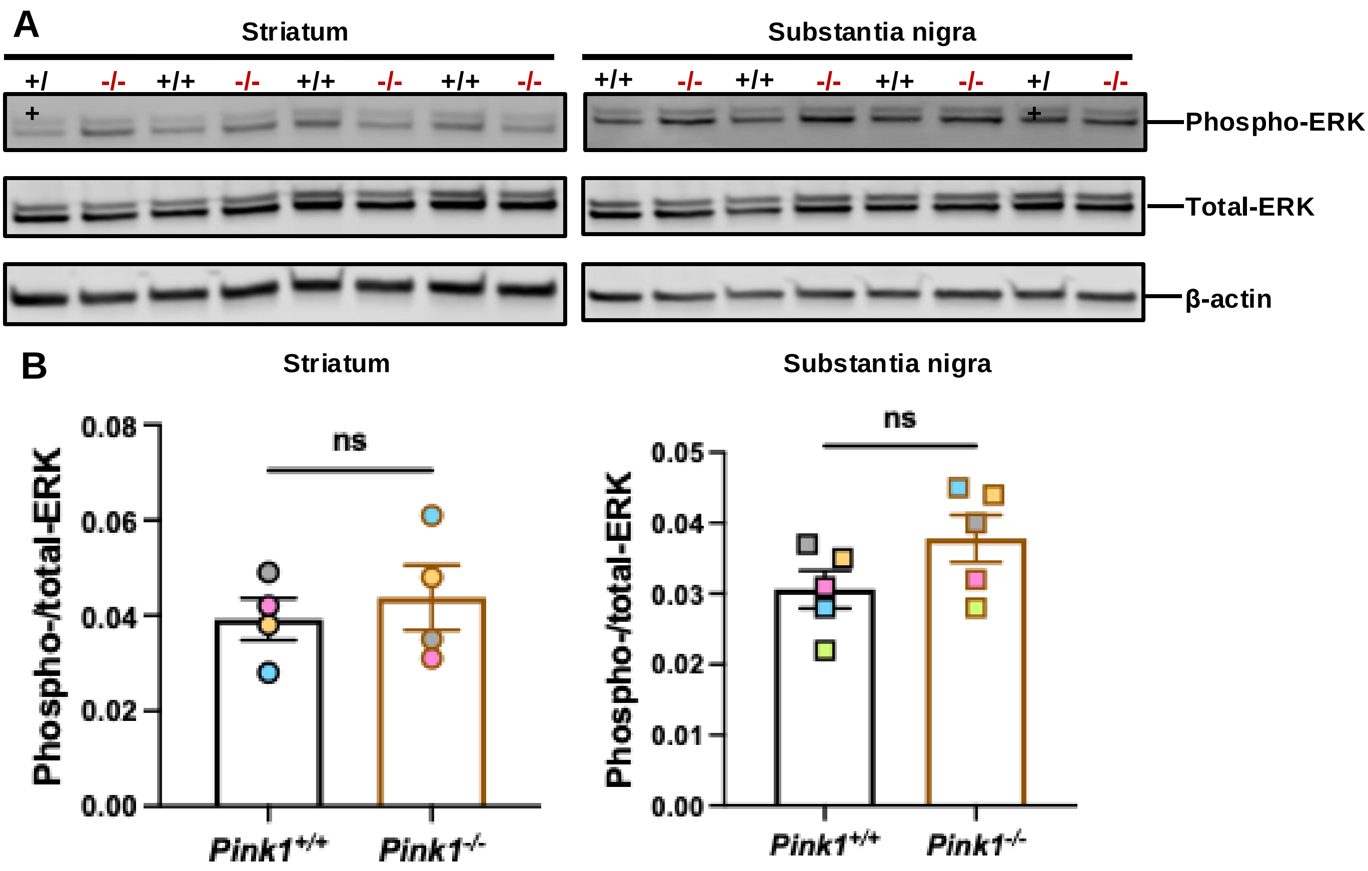


**Fig. S5 Unaltered ERK pathway activation in *Pink1*$^{-/-}$ mice. (A)** Representative immunoblots showing phospho-ERK and total ERK expression levels in the striatum (left) and substantia nigra (right) of 5-7-month-old *Pink1*$^{+/+}$ and *Pink1*$^{-/-}$ mice. **(B)** Quantification of phospho-ERK and total ERK expression in the striatum (left) and substantia nigra (right). Statistical analysis was performed using the Mann-Whitney U test.

Zhao J *et al.* Fig. S6.

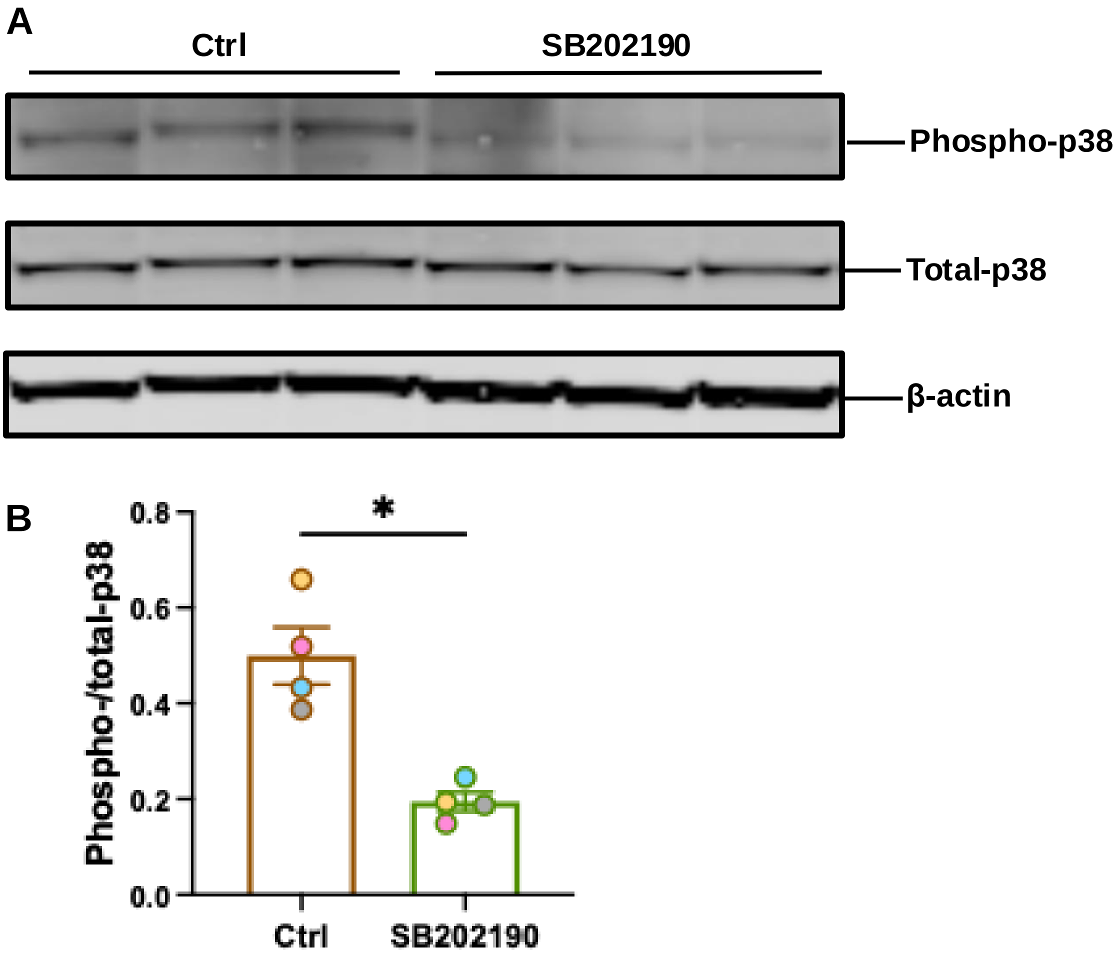


**Fig. S6 Reduced p38 phosphorylation by SB202190 treatment.** **(A)** Representative immunoblots showing phospho-p38 and total p38 expression levels before and after 50 μM SB202190 treatment. **(B)** Quantification of phospho-p38 and total p38 levels from panel A. Statistical significance was determined using the Mann-Whitney U test. $^*P < 0.05$.

**Zhao J *et al.* Fig. S7**

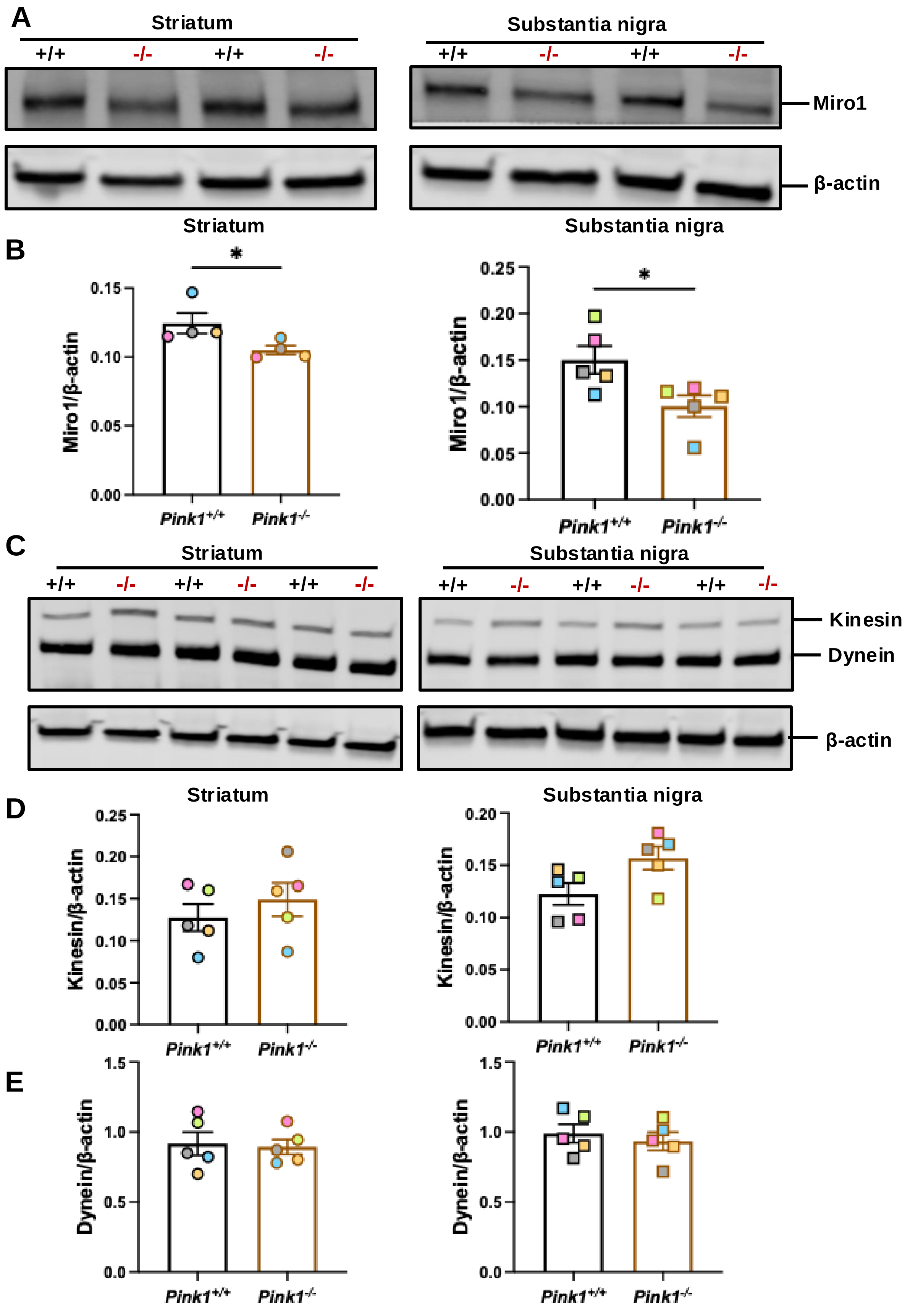

**Fig. S7 Expression levels of mitochondrial transport-associated proteins in *Pink1*$^{-/-}$ mice. (A)** Representative immunoblots showing Miro1 expression in the striatum (left) and SN (right) of 5-7-month-old *Pink1*$^{+/+}$ and *Pink1*$^{-/-}$ mice. **(B)** Quantification of Miro1 expression in the striatum (left) and substantia nigra (right) of 5-7-month-old *Pink1*$^{+/+}$ and *Pink1*$^{-/-}$ mice. Statistical significance was determined using the Mann-Whitney U test. $*P < 0.05$. **(C)** Representative immunoblots showing kinesin and dynein expression levels in the striatum of 5-7-month-old *Pink1*$^{+/+}$ and *Pink1*$^{-/-}$ mice. **(D)** Quantification of kinesin expression levels in the striatum (left) and the substantia nigra (right) of 5-7-month-old *Pink1*$^{+/+}$ and *Pink1*$^{-/-}$ mice. Statistical significance was determined using the Mann-Whitney U test. **(E)** Quantification of Dynein expression levels in the striatum (left) and the substantia nigra (right) of 5-7-month-old *Pink1*$^{+/+}$ and *Pink1*$^{-/-}$ mice. Statistical significance was determined using the Mann-Whitney U test.

**Video S1: Representative video demonstrating the established procedure for capturing mitochondrial movement along axons. Left panel: substantia nigra; Right panel: striatum.**

**Video S2: Representative video showing mitochondrial movement in *Pink1*$^{+/+}$ brain slices under normal conditions.**

**Video S3: Representative video showing mitochondrial movement in *Pink1*$^{+/+}$ brain slices after 2-hour incubation with 5 µM CGP37157.**

**Video S4: Representative video showing mitochondrial movement in *Pink1*$^{+/+}$ brain slices under normal conditions.**

**Video S5: Representative video showing mitochondrial movement in *Pink1*$^{+/+}$ brain slices after 2-hour incubation with aCSF containing 0.2% DMSO.**

**Video S6: Representative video showing mitochondrial movement in *Pink1*$^{+/+}$ brain slices after 30-minute incubation with 50 µM menadione.**

**Video S7: Representative video showing mitochondrial movement in brain slices from 7-month-old *Pink1*$^{-/-}$ mice without SB202190 treatment.**

**Video S8: Representative video showing mitochondrial movement in brain slices from 7-month-old *Pink1*$^{-/-}$ mice after treatment with 50 µM SB202190, demonstrating significant improvement in mitochondrial motility.**
